# Using AlphaFold2 to Predict the Conformations of Side Chains in Folded Proteins

**DOI:** 10.1101/2025.02.10.637534

**Authors:** Gia G. Maisuradze, Abhishek Thakur, Kisan Khatri, Allan Haldane, Ronald M. Levy

## Abstract

AlphaFold has revolutionized protein structure prediction by accurately creating 3D structures from just the amino acid sequence. However, even with extensive research validating its overall accuracy, a key question remains: Can AlphaFold predict the conformation of individual amino acid residue side chains within a folded protein? This is important for the field of molecular modeling, particularly when predicting the effects of mutations on protein stability and ligand binding. AlphaFold generates a set of atomic coordinates not just for the mutated side chain but also for potential rearrangements across the entire protein structure. In this study we investigate the ability of ColabFold, an online implementation of AlphaFold2 (AF2), to predict the conformations of residue side chains in folded proteins. We find that over a set of 10 benchmark proteins, the side chain conformation prediction error of ColabFold is on average ∼14% for χ_1_ dihedral angles, and increases to ∼48% for χ_3_ dihedral angles. The prediction error is smaller for non-polar side chains and is somewhat improved using structural templates. ColabFold demonstrates a bias towards the most prevalent rotamer states in the protein data bank (PDB), potentially limiting its ability to capture rare side chain conformations effectively. As an application of AlphaFold to explore the structural consequences of strongly cooperative mutations on side chain rearrangements, we employ a Potts sequence-based statistical energy model to perform large scale mutational scans of two proteins ABL1 and PIM1 kinase, searching for the most strongly cooperative mutational pairs, and then use ColabFold to predict the structural signatures of this cooperativity on the interacting side chains. Our results demonstrate that integration of the sequence-based Potts model with AlphaFold into a single pipeline provides a new tool that can be used to explore the fundamental relationship between protein mutations, cooperative changes in structure, and fitness.

## Introduction

To perform their functions in living organisms, most proteins must fold into a unique three-dimensional structure, or more generally, into ensembles of structures which constitute distinct functional states. These ensembles taken together characterize the protein conformational landscape. Starting from the famous experiments of Anfinsen et al. (1), not only is the question of how proteins reach their biologically active ensembles of conformations not fully answered, but also determining the three-dimensional folded structures of proteins remains one of the most challenging problems. Several experimental techniques, such as nuclear magnetic resonance (NMR) (2), X-ray crystallography (3), and cryo-electron microscopy (cryo-EM) (4), can be used to determine protein structures; however, the structures of only ∼ 0.1% (∼ 200,000 structures) of the entire protein universe have been determined. To this end, AlphaFold2 (AF2) (5), a machine-learning-based model to predict protein structures developed by DeepMind, has been groundbreaking for predicting protein structures and interactions (6). It took DeepMind only one and a half years to release structures of more than 200 million proteins predicted by AF2, i.e., structures of almost all the known proteins on the planet. Notably the 2024 Nobel Prize in Chemistry was awarded to two of the developers of AlphaFold for this achievement (https://www.nobelprize.org/prizes/chemistry/2024/summary/).

While the AF2 algorithm represents a significant advancement in protein structure prediction, one of the unresolved questions is: how accurate are these predicted structures for various computational modeling applications? Free Energy Perturbation (FEP) simulations are used to estimate the thermodynamic effects of mutations on protein stability by estimating the folding free energy difference between wild-type and mutant structures (7–13). The accuracy of these calculations depends on the accurate representation of the three-dimensional structure of the residue side chains in the folded protein (14–17). While current computational tools for predicting side chain conformations are widely used, they have not yet incorporated machine learning methods. AF2 offers an approach for side chain conformational (rotamer state) predictions by generating detailed atomic coordinates for the entire protein which captures both the local backbone and side chain structural changes at the mutation site, and potentially larger rearrangements in the protein structure.

First, it should be noted that the structure predictions by AF2 are much more accurate than by other prediction algorithms, as demonstrated at the 14th Critical Assessment of Structure Prediction (CASP14) competition (18). Second, it has recently been reported that the prediction accuracy by AF2 is often within the range of the accuracy of experimentally-derived structures of the same protein (19,20). In particular, using the root mean square deviation of C^α^ atoms, as a measure of the accuracy, the authors found that the average root mean square deviation of C^α^ atoms between different experimental structures (e.g. crystallized in a different space group) of the same protein is ∼0.6 Å, whereas it is ∼1.0 Å between experimental structures and the structures predicted by the AF2 (19). Moreover, it was shown that the AF2 models for nine small, monomeric NMR protein structures, not used in the training of AF2, have accuracies that are comparable to the experimental NMR models deposited in the PDB (20). Despite this relatively high accuracy, it was found that sometimes predictions are incompatible with experimental structure. For example, using another measure of the accuracy of the AF2 predictions, the local distance difference test (LDDT) scores, which evaluate local distance differences of all atoms in a model by comparing the inter-atomic distances of one structure to the same distances in the other structure (21) and correlate well with the confidence level of the prediction, which is quantified as the predicted-LDDT (or pLDDT), it was noted (19) that ∼ 20% of the side chains in moderate (for 70 < pLDDT < 90) to high (for pLDDT > 90) confidence residues of predicted structures had different conformations than in the experimental structure, and about 1/3 of these (7%) were incompatible with the experimental data. These findings are in general agreement with a previous study by DeepMind (22), in which 7% (for pLDDT > 90) to 30% (for 70 < pLDDT < 90) of side chains had a χ_1_ angle deviation of at least 40° from experimental structures. It was pointed out (19) that the lack of explicitly accounting for ligands, ions, covalent modifications or environmental conditions in AF2 predictions might be the reasons for some of these discrepancies.

In the present study, we performed a detailed analysis of side chain conformations, including all χ angles, for ten different proteins predicted by ColabFold (23) to determine (i) for which amino acid residues are side chain conformations most difficult to predict correctly; (ii) whether the addition of structural templates improves the accuracy of the prediction; (iii) if there are some other reasons in addition to those mentioned above that may cause these discrepancies. Here, we consider a side chain prediction is correct if it is within +/− 40° from the experimental value (17).

## Methods

ColabFold (23), a fast and user-friendly implementation of AF2, was employed to predict the structures of ten different proteins, including, six α+β proteins [staph nuclease (PDB ID: 1EY0 (24)), T4 lysozymes (PDB IDs: 2LZM (25) and 1L63 (26)), ABL-1 kinase (PDB ID: 2V7A (27)), PIM-1 kinase (PDB ID: 1XWS (28)), and antifreeze protein (PDB ID: 3VN3 (29))], two β sheet proteins [choristoneura fumiferana (spruce budworm) antifreeze protein (PDB ID: 1L0S (30)) and unknown function protein (BF3112) from bacteroides fragilis NCTC 9343 (PDB ID: 3MSW (31))], and two α helical proteins [myoglobin (PDB ID: 3V2V (32)) and the transmembrane protein TMHC2_E (PDB ID: 6B87 (33))]. By replacing AF2’s homology search with the 40 to 60-fold faster, Many-against-Many sequence searching (MMseqs2 (34,35)), ColabFold significantly accelerates predictions of protein structures and complexes. The structure predictions on ColabFold were performed with– and without structural templates. The proteins selected in this study are from diverse families, and differ from each other by size and biological function. Predicted structures have been validated by comparing them to experimental PDB structures.

ColabFold allows users to customize the protein structure prediction. There are several important parameters, including particularly the depth of the multiple sequence alignment (MSA), and the use of structural templates, that can be changed. Supplying a structural template, pushes ColabFold prediction to resemble the provided structure, although the extent to which the prediction is biased by the template and the accuracy of the prediction depends on the MSA depth, as well. In this study, we used the target proteins themselves as the structural templates, which enabled us to investigate the contributions from the structural template and the MSA in the prediction accuracy.

## Results and Discussion

### Application of structural templates for side-chain structure prediction and performance by residue type

We have performed structure predictions by ColabFold with and without structural templates for ten different proteins to elucidate the prediction accuracy of side chain conformations in the folded protein: (a) when only the MSA and the sequence of the target protein are provided; and (b) to determine how the prediction accuracy changes with the addition of a structural template.

The results illustrated in Figure 1 show that the prediction accuracy for side chain conformations with and without templates, on average, is highest for χ_1_ angles and decreases with the increase of χ index (the average errors for χ_1_ angles are 12% and 17% with and without templates, respectively, and increase to 47% and 50% with and without templates on average, respectively, for χ_3_ angles. The χ_4_ angles are an exception because of their existence only in two amino acids – Arginine and Lysine). Moreover, the prediction accuracy of χ depends on the depth of the MSA and the secondary structure. ColabFold tends to predict side chain χ_1_ dihedral angles for proteins with α + β structures more accurately than ones for proteins with α-helical or β-strand only structures.

**Figure 1.**
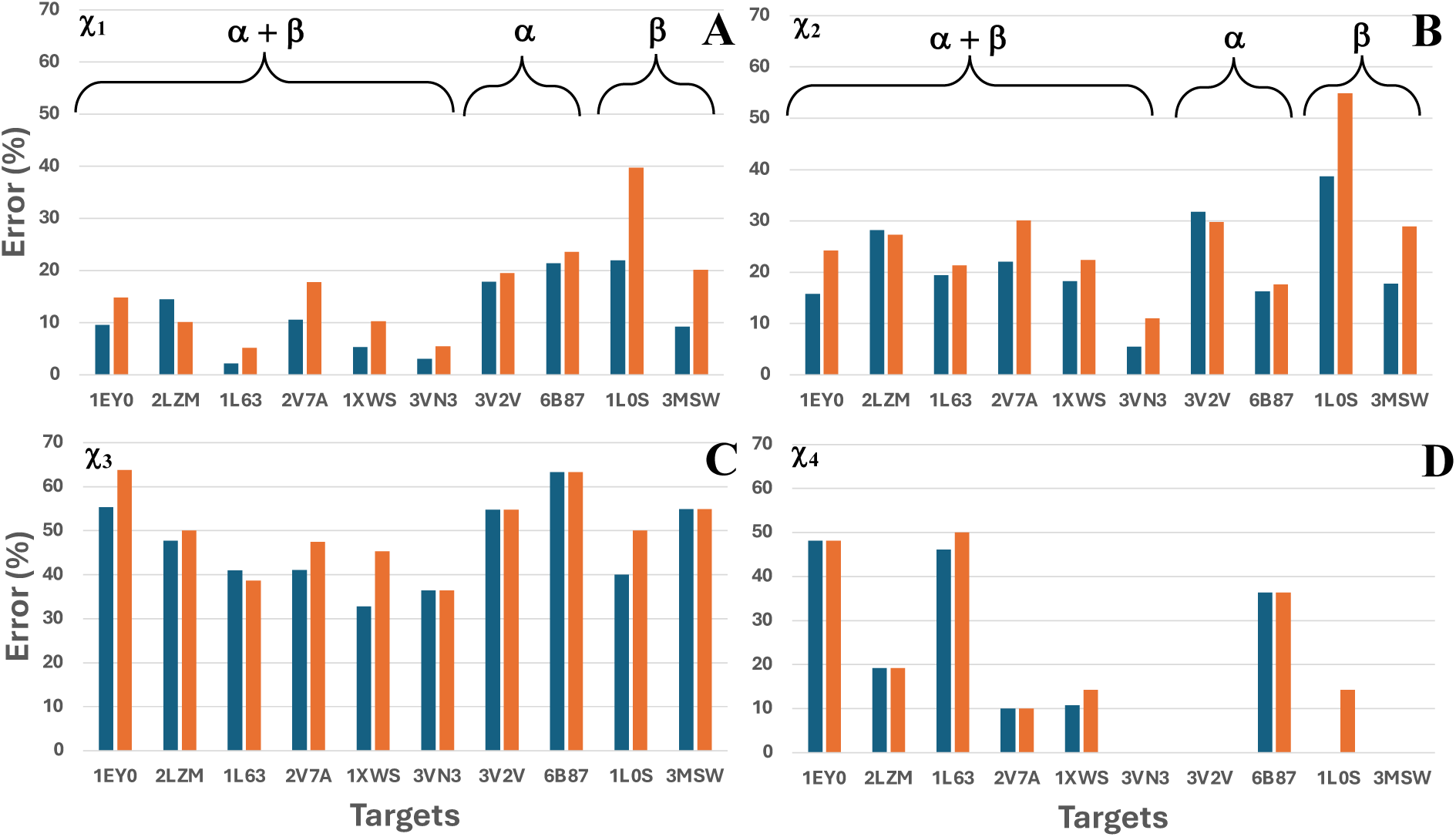
The percentages of incorrectly predicted χ_1_ (panel A), χ_2_ (panel B), χ_3_ (panel C), and χ_4_ (panel D) angles, in predicted structures with (blue bars) and without (orange bars) templates, determined by the formula: 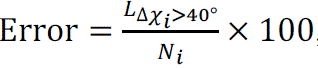, where 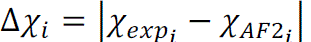, *i =* 1, 2, 3 and 4, *L_i_* is number of predicted *χ_i_* angles, values of which differ from their experimental counterparts by more than 40°, *N_i_* corresponds to the number of all *χ_i_* angles.

The utilization of structural templates in protein structure prediction by ColabFold results in an enhancement in the accuracy of the predicted rotamer states of side chains in the ten benchmark proteins (∼ 15% averaged over all side chain torsions) (Figure 1). The addition of templates has a significant effect on the χ_1_ conformations (improvement in the accuracy of ∼31% on average using templates) but very little effect on the accuracy of χ_3_ predictions. There is one protein (2LZM) among ten studied proteins, for which the χ_1_ and χ_2_ angles exhibit an opposite trend; the side chain prediction accuracy is better without the structural template (Figure 1). The accuracy of the ColabFold structure prediction strongly depends on the MSA. A diverse and deep MSA, with thousands of sequences in the alignment, helps ColabFold to identify coevolutionary signals, and when these coevolutionary signals are strong ColabFold tends to ignore template structures. 2LZM has a deep MSA (> 3,500 sequences), and might be one of the reasons for the inconsistency. Secondly, the depth of the MSA and the inclusion of a structural template mainly affect the accuracy of the backbone conformations of predicted structures, whereas the accuracy of side chain conformations of some amino acids does not always rely on these parameters. The factors playing a role in inaccurate prediction of side chain conformations are investigated below in this section and following sections.

Our results in Figure 1 illustrate that incorporating a structural template has a significantly greater effect improving the accuracy of proteins assembled from β sheets (e.g., 1.8 times in 1L0S, and 2.2 times in 3MSW), which suggests a potential advantage of template-based approaches for β-sheet prediction. β-sheets, characterized by tightly packed, antiparallel strands connected by hydrogen bonds, can exhibit significant structural variability, which is usually not the case for α-helical proteins. Moreover, shallow MSAs are another reason for the noticeable improvement when template structures are used; in particular, 1L0S and 3MSW have only ∼ 40 and ∼ 400 sequences, respectively, in the alignments.

In Figure 2, we decompose the overall performance of ColabFold for predicting side chain dihedral angles in folded proteins by residue type. The percentages of incorrectly predicted χ angles for the most polar amino acids are higher than for non-polar amino acids in both with and without template predicted structures. It should be noted that ColabFold with and without templates, predicts correctly all χ angles for two hydrophobic aromatic amino acids tryptophan and tyrosine (the total number of TYR + TRP residues in the ten target proteins is 93). Furthermore, Figure 2 reveals a general trend of lower prediction errors for χ_1_ angles compared to χ_2_ and χ_3_ (as in Figure 1, χ_4_ angles are an exception due to their existence only in two amino acids) across all residue types. This is related to the inherent greater flexibility of χ angles closer to the side chain terminus compared to the more constrained χ_1_ which are subject to restriction due to steric hindrance between the ψ side-chain atoms and the main chain. In all cases the AF2 predictions with templates have lower error than without templates. Moreover, Figure 2 highlights a significant improvement (∼ 20% error reduction) in the prediction of the χ_1_ dihedral angles for Cysteine (Cys) residues upon template addition. This observation suggests that AF2 might benefit from modifications of the objective function specifically focused on accurately predicting the χ_1_ rotamer state for Cys residues (36). It is important to mention that only 21 cysteine residues are observed in the ten studied proteins (five of them do not have cysteine residues at all). All errors in χ_1_ dihedral angles are found in 1L0S, which contains the most cysteines (eight residues). The structure of 1L0S reveals a left-handed β-helical structure consisting of 15 amino acid loops. Each helix loop has three sides (β strands) and Ser/Thr/Cys stacking within the interior of the protein at the corners of the β helix strengthened by four interstrand disulfide bonds formed by cysteines (30). All four interstrand disulfide bonds and eight χ_1_ dihedral angles of cysteines are correctly predicted in ColabFold predicted structure with template. The ColabFold predicted structure without template reveals only two disulfide bonds formed by cysteine residue pairs, although these pairs are not the same as in the 1L0S PDB structure and ColabFold predicted structure with template; plus, four χ_1_ dihedral angles of cysteines are incorrectly predicted. This finding indicates that not only side chains but also backbone of 1L0S is predicted poorly by ColabFold without template (see sub-section “Correlation between pLDDT, backbone and side-chain angles” below). Notable improvement was also observed in the prediction of χ_3_ angles for Methionine (Met).

**Figure 2.**
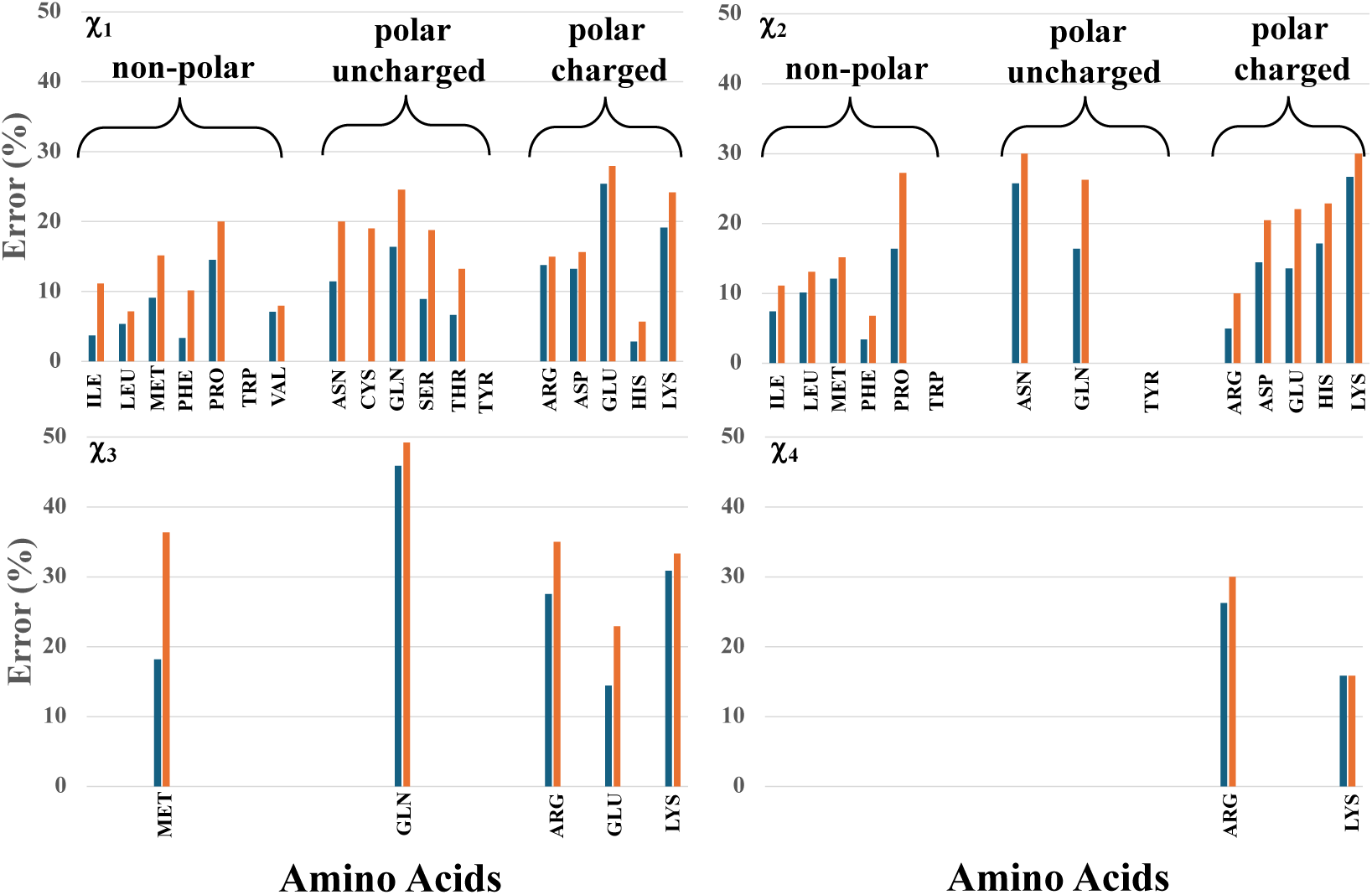
The same as in Fig. 1 but for amino acids.

### AlphaFold2 bias towards dominant rotamer states: Implications for rare conformations

Our analysis of AF2’s rotamer state predictions reveals a potential bias towards the most frequently observed rotamer conformations in the PDB (dominant states). This bias suggests the difficulties that AF2 may have predicting rare rotamer states, even when these rare conformations are present in the experimental crystal structure provided as a template. By “rare,” we refer to conformations observed in a very small percentage (< 1%) of protein data bank entries, according to the Schrödinger knowledge base (Schrödinger suite release 2023-2) that contains the rotamer state populations for each amino acid based on an analysis of the PDB (37,38). More precisely, out of a total 1453 side chain rotamer states, predicted in this work, 8.2% of the experimental side chain conformations are rare. For visual illustration, we superposed experimental and AF2 predicted structures of staph nuclease (1EY0) (Figure 3A) and T4 lysozyme (2LZM) (Figure 3B), and selected two residues (LYS-49 in 1EY0 and LYS-16 in 2LZM) with rare and PDB dominant rotamer states in experimental and predicted structures, respectively. As shown in Figure 3, side chains of both residues have enough room to adopt various conformations, however, there are some differences. In particular, in 1EY0, LYS-49 is in a loop in both the experimental and AF2 predicted structures, which have a large deviation from each other [backbone root mean square distance (RMSD) at residue 49 is 1.74 Å], that permits the formation of a salt bridge between the side chains of LYS-49 (orange) and GLU-52 in the AF2 predicted structure, not observed in the PDB structure. In 2LZM, LYS-16 is embedded in different secondary structures (β-sheet in experimental structure and loop in AF2 predicted structure), however, these secondary structures are only slightly deviated from each other (backbone RMSD at residue 16 is 0.59 Å), and hence, probably do not play a role in the discrepancy between the predicted and experimental rotamer states of LYS-16. It should be noted that ∼ 55% and ∼ 42% of errors for χ_1_ angles and for rotamer states, respectively, in all ten studied proteins come from the loop regions. Also, rotamer states of only ∼ 1/3 of loop regions are incorrectly predicted by AF2.

**Figure 3.**
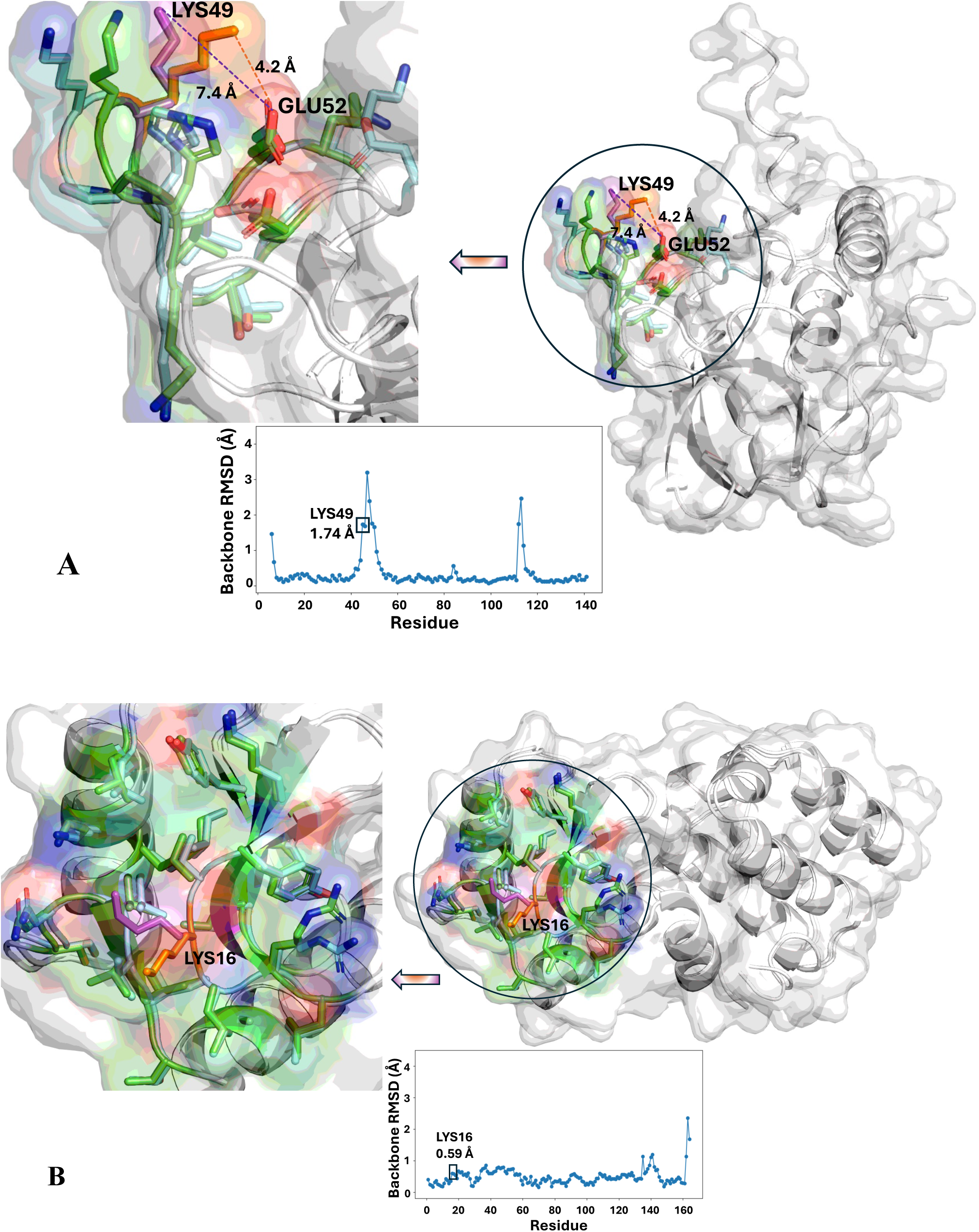
Superposed experimental and AF2 predicted structures along with RMSDs of C^α^ atoms between these two structures for staph nuclease (panel A) and T4 lysozyme (panel B).

Using the Schrödinger database we have assigned the closest rotamer state in the PDB-derived database for each residue over the entire sequence for the AF2 predicted and the experimentally determined protein structure and plotted those values in the X and Y axis of Figure 4, respectively. Because there are a large number of points in Figure 4 which sometimes overlap and also differences in the distribution of rotamer state populations for amino acids in the Schrödinger knowledge base (Table S1 shows the percentages of five most dominant rotamer states of each amino acid), in Supporting Material we additionally (i) plotted the results separately for each amino acid (Figure S1); (ii) calculated the ratios between the PDB derived database percentages for each residue of ten experimental and AF2 predicted structures (Figure S2); (iii) computed probability distribution functions (PDFs) of rotamer states of each amino acid obtained from ten experimental and AF2 predicted structures along the populations of rotamer states in protein data base (Figure S3); and (iv) computed how frequently each amino acid in experimental and AF2 predicted structures adopt more dominant rotamer state from protein data base (Figure S4).

**Figure 4.**
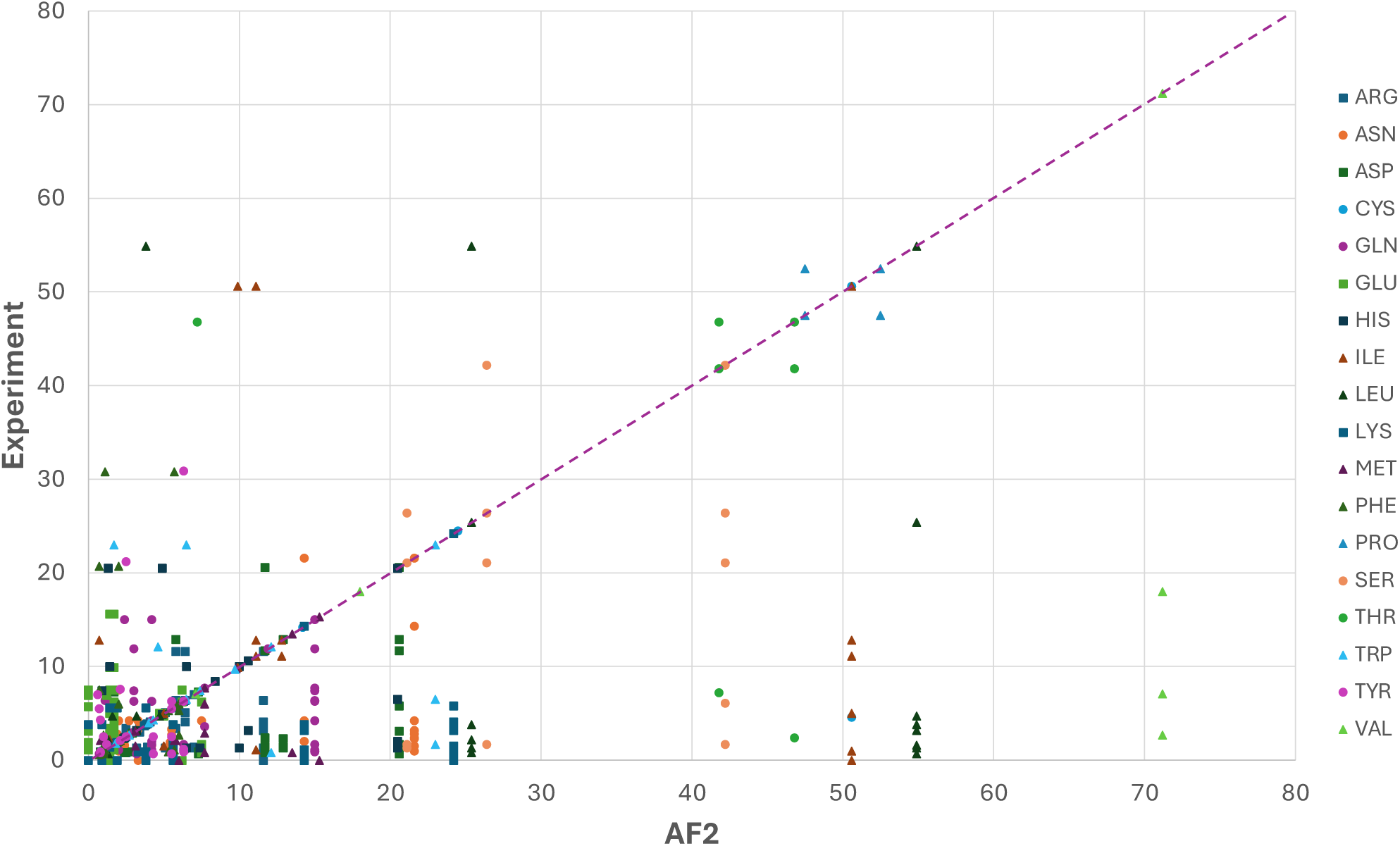
Comparison of rotamer states of each amino acid [non-polar (triangles), polar uncharged (circles), polar charged (squares)] of experimental and AF2 predicted (with template) structures in terms of the probability (in %) of the closest rotamer state in the protein database.

The results show that rotamer states of most of the amino acids, except for Glu, Phe, and Tyr, in the AF2 predicted structures tend to adopt the dominant conformations from the protein data base. That is, the majority of triangles, circles, and squares in Figure 4 are located below the diagonal indicating that AF2 frequently predicts the most populated states whereas rotamer states of these amino acids obtained from experimental structures of the ten studied proteins are either in less populated states or rarely observed in the PDB database. It is not surprising that the most easily visible residues, in Figure 4, showing a bias towards the most frequently observed conformations in protein databases in AF2 predicted structures are Leu, Ile and Val, which have much higher percentages of dominant rotamer states than others (see Table S1). Further, as illustrated in Figure S4, (i) the rotamer states of polar amino acids Arg, Asn, Asp, Gln, and Lys in AF2 predicted structures exhibit a strong inclination towards more dominant rotamer states in protein data base than their counterparts in experimental structures; (ii) Glu, Phe and Tyr show opposite results, i.e., rotamer states of these amino acids in experimental structures exhibit a strong tendency to adopt more dominant rotamer states in protein data base than their counterparts in AF2 predicted structures; (iii) the rotamer states of Pro and Trp in experimental and AF2 predicted structures exhibit an equal propensity towards the dominant rotamer states in protein data base; (iv) the rotamer states of rest of the amino acids in AF2 predicted structures are more inclined towards the dominant rotamer states in protein data base than their counterparts in experimental structures, although in majority cases rotamer states of these amino acids in AF2 and experimental structures do not differ from each other and fall into the same category defined by protein data base. These results suggest training bias in the AF2 model that corresponds to a potential limitation in AF2’s ability to handle low population rotamer states when predicting the protein structure.

It is well known that “rare” rotamer states might represent local minima in the protein’s energy landscape (16). Hence, without MD refinement that accounts for various interactions, such as hydrogen bonding, steric clashes, and solvation effects, capturing these subtle energy minima can be difficult for the AF2 algorithm (39). This also suggests that AF2’s internal sampling procedures might not be sufficiently comprehensive to explore the full conformational space, especially for regions with high flexibility or the presence of rare rotamers.

### Spatial clustering of rotamer prediction errors adjacent to Alanine and Glycine residues

One of the most intriguing observations from our analysis of multiple ColabFold-predicted PDB structures is the spatial clustering of rotamer prediction errors. We observed a frequent trend where prediction errors tend to occur next to Alanine (Ala) or Glycine (Gly) residues in the protein sequence. In particular, ∼20% of all errors in the range of 40° – 100°, found in predicted structures for χ_1_ angles occur in residues next to Ala or Gly. Moreover, the error rate increases to ∼40% for errors Δχ_1_ ≥ 100°.

One possible explanation lies in backbone flexibility. Both Ala and Gly lack bulky side chains, leading to a more flexible backbone compared to other residues. This inherent flexibility can introduce conformational heterogeneity, particularly in neighboring peptide planes.

To explore this problem, we computed root mean square distances between the residues of experimental and AF2 predicted structures for ten studied proteins, and found that of the side chain prediction χ_1_ errors adjacent to Gly and Ala residues ∼ 66% might be associated with a lack of packing constraints due to the small size of Ala and Gly, and the remainder (∼34%) might be associated with backbone prediction errors, respectively. For visual illustration of both mechanisms, we selected one of the proteins (2LZM), which exhibits large errors (> 40°) in rotamer states at eight residues next to Ala and Gly. Two curves are depicted in Figure 5: the blue curve corresponds to errors between χ_1_ angles obtained from experimental and AF2 predicted structures, and the orange curve corresponds to backbone RMSDs. Because Ala and Gly do not have χ angles, the blue curve looks like a piecewise function, in which the “breaks” correspond to Ala and Gly residues. Based on the behavior of the RMSD over the sequence, the threshold at 0.5Å was determined indicating which mechanism may cause a large error in rotamer state. In other words, if the backbone RMSD > 0.5Å, a large discrepancy in rotamer state might be caused by backbone prediction errors, and if RMSD ≤ 0.5Å, a large error in rotamer state might be induced by lack of packing constraints. All eight residues are numbered with red (RMSD > 0.5Å) and black (RMSD ≤ 0.5Å) colors depending which mechanism may cause the large discrepancies in χ_1_ angles. Also, two inserts are included in Figure 5 illustrating the structural differences caused by these two mechanisms. The insert for Lys-48 shows the discrepancy between the backbones of experimental and AF2 predicted structures, which may consequently originate the difference in rotamer states, however, due to a large free space in the neighborhood of Lys-48 the tendency of AF2 to adopt the more dominant rotamer states in protein data base could easily cause this discrepancy in χ_1_ angles. The rotamer state of Lys-48 in the experimental structure has a frequency of 0.9% in the PDB database, while the AF2-predicted rotamer state has a frequency of 5.6%. The insert for Lys-147 demonstrates that there is a large free space in the vicinity of Lys-147 allowing side chains to adopt different conformations. While the small size of Ala may be the reason for the discrepancy in χ_1_ angles in Lys-147, it is possible its side chain could adopt different conformations with having other neighbor residues, as well. The main reason for the discrepancy may again be due to AF2’s bias for dominant rotamer states, because the rotamer state of Lys-147 in the experimental structure has frequency very close to zero (very rare) in the protein data base, while it is 5.6% in AF2 predicted structure.

**Figure 5.**
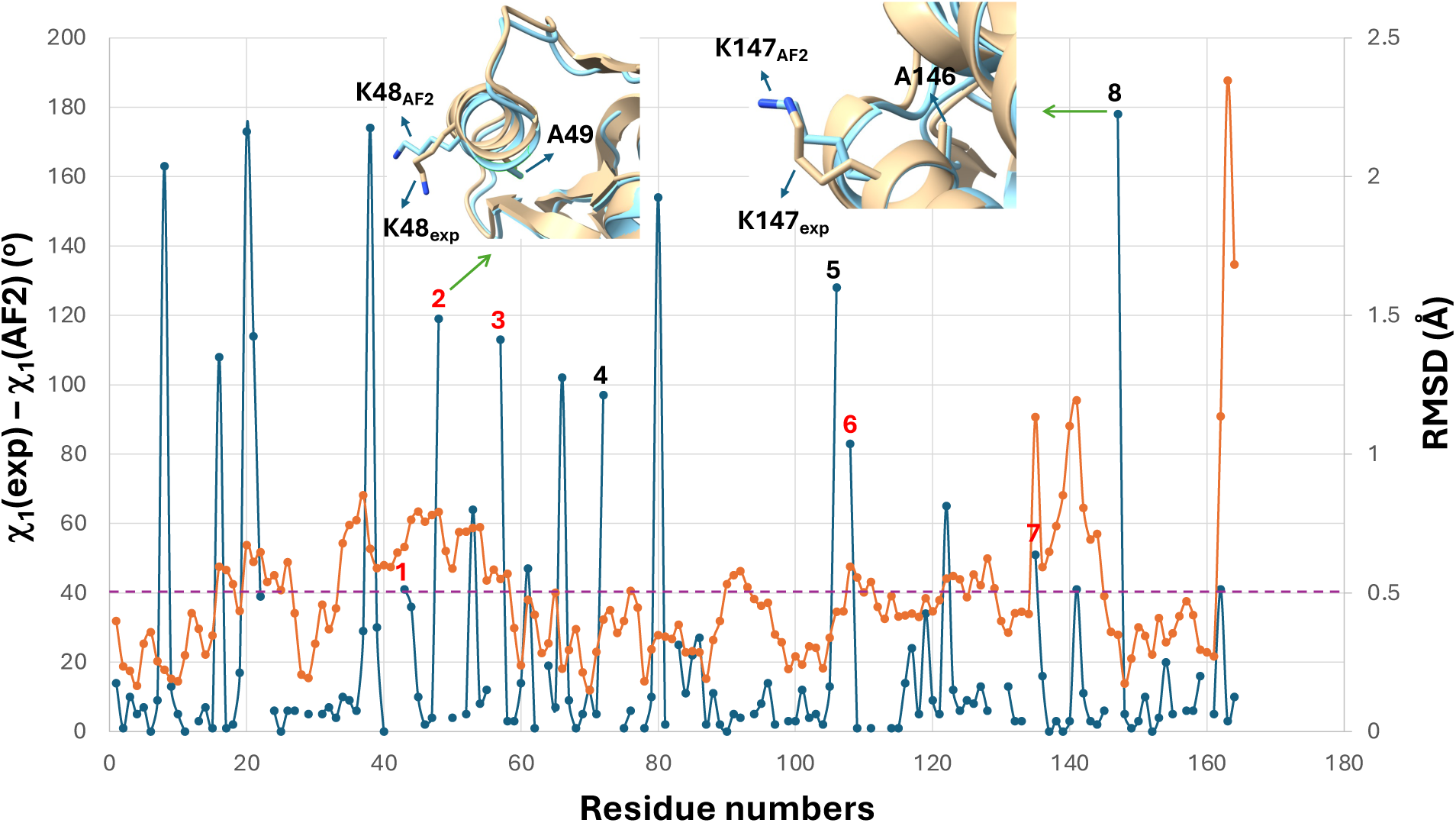
Absolute differences between χ_1_ values (blue curve) and RMSDs (orange curve) of experimental and AF2 predicted structures with template for 2LZM. Red color numbers indicate large errors in rotamer states that might be caused by backbone prediction errors, black color numbers indicate large errors in rotamer states that might be induced by lack of packing constraints. Inserts with experimental (gold) and AF2 predicted (cyan) structures illustrate the differences between side chain conformations, in particular, for Lys-48 and Lys-147, caused by these two mechanisms. Purple dashed line represents the threshold for χ_1_ angles (40°) and RMSD (0.5Å).

To understand whether discrepancies between backbones of experimental and AF2 predicted structures are correlated with errors of side chain rotamer states, we computed (i) the Pearson correlation coefficients between the RMSD and Δχ_1_ values; (ii) average RMSD values for all Ala and Gly residues, separately, and for entire sequences without Ala and Gly residues; and also, for all residues with large rotamer state errors (Δχ_1_ > 40°) (see Table S2). The results show that (i) there is either very weak or no correlation between RMSD and Δχ_1_ values; (ii) average RMSD values are changing (“oscillating”) from protein to protein. However, as was expected, average RMSD values for residues with large rotamer state errors are greater than average RMSD values for Ala, Gly and for rest of sequences in most proteins. It should be noted that large discrepancies (> 0.5Å) between backbones of experimental and AF2 predicted structures are mainly found in loops and ends (only two proteins, 3V2V and 6B87, exhibit opposite results) (see Table S2). Therefore, if loops contain Ala and Gly residues, average backbone error at Ala and Gly residues may increase, which is the reason of significantly large average RMSD values for Ala and Gly in proteins 3MSW and 2V7A, respectively.

Based on these results, the driving force causing the discrepancies between rotamer states of residues of experimental and AF2 predicted structures is the inclination of AF2 to adopt more dominant rotamer states from protein data base. The lack of packing constraints caused by a small size of Ala and Gly or increased backbone flexibility can play a role in the discrepancies between rotamer states, although it may not be as significant as the former.

### Correlation between pLDDT, backbone and side-chain dihedral angles

As stated in the Introduction, the accuracy of the AF2 predictions is evaluated by the LDDT scores that correlate well with the confidence level of the prediction (pLDDT). It has been shown that a pLDDT score greater than 90 is taken as the benchmark for very high confidence and a pLDDT score greater than 70 is considered to be a moderate-to-high confidence structure and corresponds to a correct backbone prediction, and a pLDDT < 50 indicates unreliable predictions (19). Moreover, it was observed that AF2 can predict side-chains (c_1_ angles) with high accuracy when the backbone prediction is high confidence (pLDDT >90) (22). It is of interest to know how the pLDDT scores are correlated with the prediction accuracy of backbone and side-chain angles. Among the ten studied proteins we selected three proteins which illustrated different behavior. In particular, (i) the predicted structures of staph nuclease (PDB ID: 1EY0) with and without template have very high pLDDT (> 90) along the entire sequence except for one loop region (43-53, 62 < pLDDT < 85) (Fig. S5A); (ii) the predicted structures of T4 lysozyme (PDB ID: 1L63) with and without template exhibit very high (> 90) pLDDT score along almost entire sequence (Fig. S5B); (iii) the predicted structure of choristoneura fumiferana (spruce budworm) antifreeze protein (PDB ID: 1L0S) with template shows very high (> 90) pLDDT score along the entire sequence, whereas the predicted structure without template illustrates moderate and below moderate confidence (Fig. S5C). Figure S5 shows moderate negative (for 1EY0 and 1L0S) and weak negative (for 1L63) correlations between pLDDT and backbone angles (ψ,ϕ) and very weak negative correlation between pLDDT and χ_1_ angles for all three proteins. There is no correlation observed between pLDDT and χ_2_, χ_3_, χ_4_ angles, consequently no correlation between (ψ,ϕ) and χ_2_, χ_3_, χ_4_ angles.

### Prediction accuracy of buried vs solvent exposed side chains

To elucidate how the solvent accessibility of each residue changes in the AF2 predicted structures compared with the experimental PDB structures, we computed the solvent accessible surface areas (SASAs), which is considered to be an important factor in protein folding and stability, of each residue in all ten studied proteins. Figure S6 illustrates the probability distribution functions of percentages of solvent accessibility areas of each amino acid computed from experimental and predicted structures (with and without template). As was expected, most of the non-polar residues are buried in protein except for proline, which despite its non-polar nature is not hydrophobic (40), consequently it is more solvent exposed than other non-polar residues. The differences between the results of experimental and both predicted structures are not significant, which indicate that side-chain conformations of non-polar residues are predicted correctly in terms of solvent accessibility.

The residues with uncharged polar and charged polar side chains, in general, are more exposed to the solvent than non-polar residues (Figure S6), however, side chains of some polar residues either are buried in protein (cysteine and tyrosine) or exhibiting no preference to be solvent exposed (histidine). Although the cysteine side chain is polar, it is considered hydrophobic based on the observation that it is often found in the interior of proteins largely due to its ability to form disulfide bonds. Moreover, the tyrosine is sometimes considered as a polar amino acid because its side chain has OH group, however, it also has a nonpolar benzene ring, and tyrosine is generally classified as a hydrophobic amino acid. Although histidine is a polar, positively charged amino acid, its physical properties depend very much on pH. In particular, the imidazole group of histidine is the only amino acid side chain affected by the range of physiological pH values; i.e., at pH 5.0 the group is positively charged, polar, and hydrophilic, whereas at pH 7.4 it is neutral, apolar, and hydrophobic (41). Therefore, a strong dependence of physical properties of histidine on pH might be a reason for broad distribution of percentages of SASA.

As in non-polar residues, the differences between the results of experimental and both predicted structures with and without templates for polar uncharged and polar charged residues, in general, are not significant.

It should be noted that PDFs for some polar uncharged (threonine and serine) and polar charged (aspartic acid and glutamic acid) amino acids do not exhibit strong propensity to be solvent exposed. This behavior of amino acids is in harmony with the hydropathy index defined by Kyte and Doolittle (42). Based on this work, (i) amino acids with a hydropathy index equal to or more than 1.8 are defined as hydrophobic; (ii) amino acids with a hydropathy index equal to or less than –3.3 are defined as hydrophilic; (iii) amino acids with a hydropathy index less than 1.8 and more than –3.3 are defined as neutral. Hydropathy indices for threonine and serine are –0.7 and –0.8, respectively, which indicate that both amino acids are neutral, consequently, it is not surprising that they do not exhibit strong propensity to be solvent exposed. Hydropathy index for both aspartic acid and glutamic acid is –3.5, which indicates that both amino acids are hydrophilic, however, their closeness to the threshold value (–3.3) might be a reason for a weak propensity to be solvent exposed.

### Structural signatures of strongly cooperative double mutations are discovered by combining Potts sequence co-variation models with AlphaFold

There is a long history of using co-evolutionary information encoded in protein MSAs to probe the relationship between protein structure, function, and fitness (43–48). Potts Hamiltonian models provide a simple but powerful framework to capture the sequence co-variation patterns in MSAs using a machine learning approach, which share some characteristics with transformers, the machine learning algorithm employed by ColabFold (23). Implicitly, ColabFold is exploiting information about correlated mutations in its predictions of the sequence dependence of the most stable three-dimensional structure(s) of a protein. Furthermore, being able to predict non-additive effects of pairs of mutations is important for understanding how mutations alter protein stability, protein-protein interactions, ligand binding affinity, and many other sequence-structure properties that impact protein function. We have used Potts models extensively to study the fitness and conformational free energy landscapes of kinase family proteins (43,49–52), and to investigate the effects of mutations on the acquisition of drug resistance in HIV protein targets (43,53–56). The non-linear effects of pairs of cooperative mutations on protein fitness can be estimated at scale for large numbers of mutations using the Potts model for mutational scanning on many sequences in a protein family, whereby Potts double mutant cooperativity ΔΔ*E* is calculated for all pairs of mutations in each sequence. Using Potts model mutational scanning as a preprocessing step to select mutation pairs that are predicted to be strongly coupled, followed by AlphaFold structure predictions of the corresponding sequences, is a way to identify changes in the rotamer states of side chains that may provide a possible structural explanation for the strong cooperativity indicated by the Potts model sequence-based calculations. In this section we illustrate how such modeling might proceed.

The Potts model can be used to estimate the cooperative effect of a pair of mutations on protein fitness through the calculation of ΔΔ*E*:

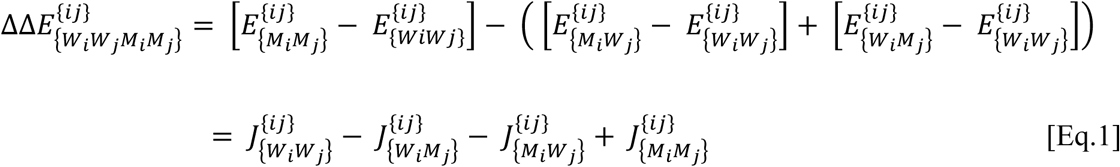

The first term in square brackets is the relative log probability of the double mutation (M*_i_*, M*_j_*) at positions *i* and *j* relative to the reference residues (W*_i_*,W*_j_*), while the second and third terms in square brackets are the corresponding log relative probabilities of the two single mutations. The field contributions to each of the terms cancel when combined in the ΔΔ*E* formula. ΔΔ*E* then depends on the four couplings between the residues pairs (W*_i_*,W*_j_*), (M*_i_*, M*_j_*), (W*_i_*, M*_j_*) and (M*_i_*, W*_j_*) at positions *i* and *j*. Importantly, ΔΔ*E* is independent of the gauge of the Potts statistical energy function. The non-linear cooperativity is a measure of how much more favorable (or unfavorable) the fitness of the double mutation is than the sum of the effects of the individual mutations on the fitness. See the Appendix for a detailed description of the Potts Hamiltonian Model and the derivation of [Eq.1].

We have used a Potts model of Kinase Family proteins to compute a histogram of ΔΔ*E* values for all pairs of mutations in ABL1 and PIM1, the results are shown in Figure 6 as ΔΔ*E* histogram plots. We found that less than 1% of the total double mutations (0.78% for ABL1 and 0.84% for PIM1) have the ΔΔ*E* values greater than 1.5, indicating that only a small fraction of mutation pairs are strongly cooperative. Among these, the 10 most positively cooperative mutation pairs appearing in the tail of the ΔΔE distribution are labelled in Figure 6.

**Figure 6.**
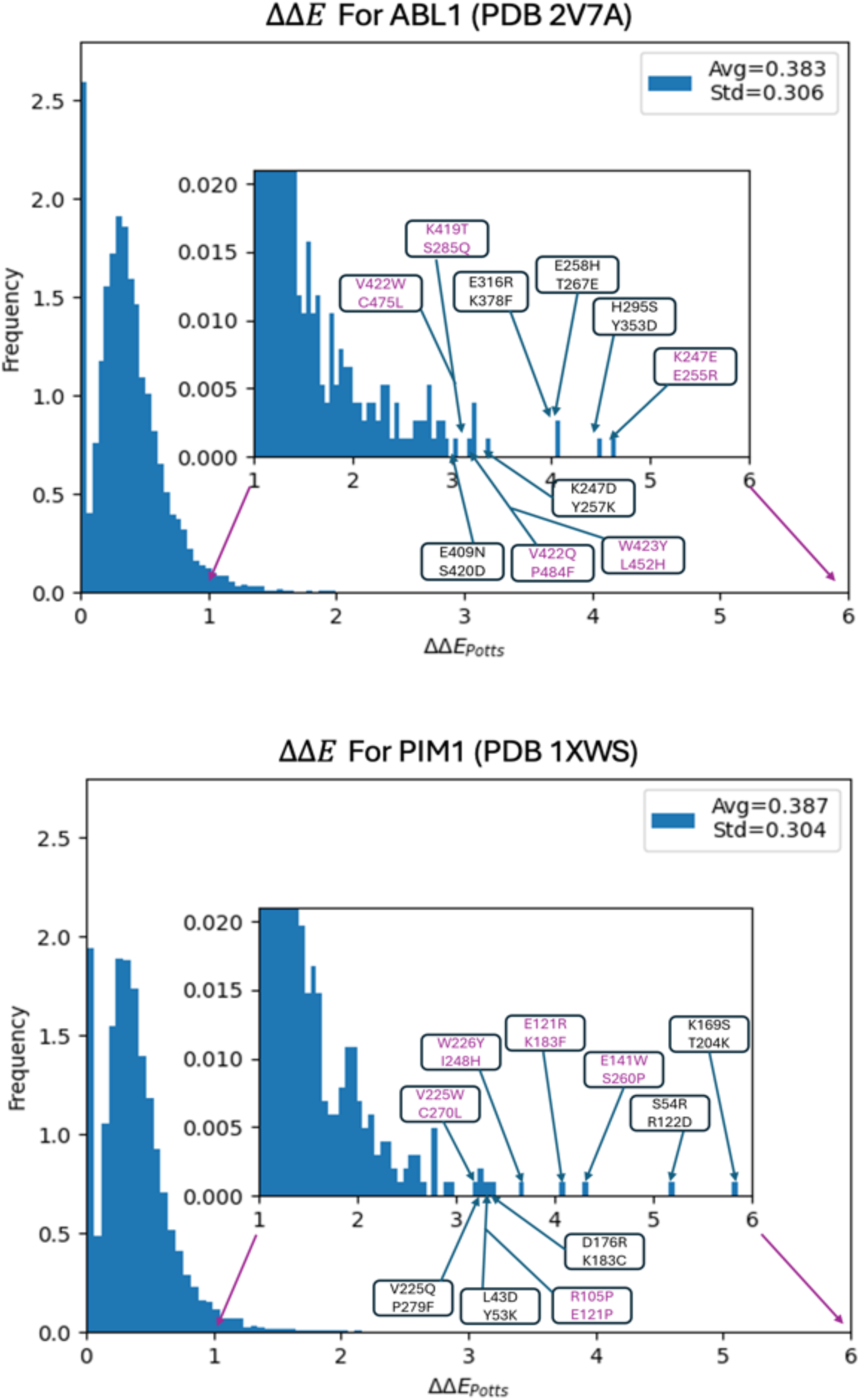
ΔΔ*E* Histogram Plots for double mutant cycle values for ABL1 and PIM1. The annotated mutant pairs represent the top 10 most positively cooperative double mutant pairs, with black indicating pairs that show the direct cooperative effect, while pink representing those whose cooperativity can be explained in an alternate way (see text).

The simplest structural signature of mutational cooperativity involving mutations at positions *i* and *j* in the sequence, is for one of the mutated residues to perturb the rotamer state of the other mutation. In other words, the rotamer state of residue M*_i_* changes depending on whether the residue at position *j* is W*_j_* or M*_j_*. To test this, three AlphaFold structure predictions are required for each mutation pair: (M*_i_*,W*_j_*), (W*_i_*,M*_j_*), and (M*_i_*,M*_j_*). We have performed this analysis for the ten mutation pairs that are predicted by the Potts model to be most strongly cooperative for both ABL1 and PIM1. For 5 of the 10 most cooperative mutation pairs predicted by the Potts model for ABL1 and PIM1, the AlphaFold structural predictions show the most direct structural signature of double mutant cooperativity (indicated by the black residue labels in the histogram, Fig. 6); AlphaFold predicts that the rotamer state of one mutant residue perturbs the rotamer state of the other mutant residue. These five cooperative mutation pairs which serve as an example, can mostly be explained as resulting from the maintenance of a charge-charge interaction involving the double mutant residues (e.g. ABL1: K247D/Y257K, and PIM1: L43D/Y53K), or by charge swapping (e.g. PIM1: K169S/T204K).

The cooperativity of several of the remaining 5 out of 10 pairs can also be accounted for (represented by pink in the histogram). In the case of the most cooperative pair K247E/E255R (ABL1), we observe the single mutant K247E in the wild-type sequence causes a rotamer change in the wild-type residue E255, supporting a cooperative effect without reference to the interactions between the double-mutant side chain rotamer states described above and listed in Table 1. It is also possible for mutation pairs to be cooperative without any rotamer state change, such as in K419T/S485Q (ABL1). The single mutant AF2 predictions show electrostatically repulsive groups in proximity while the wildtype and double mutant have electrostatically attractive groups in proximity. Even though the rotamers are locked in place by their environment, the attractive pairings of the double mutant stabilize the protein and improve folding. Others of the remaining pairs can be similarly rationalized, for instance in the double mutant W423Y/L452H (ABL1) the hydrophobically attractive pair WL is replaced by the electrostatically attractive mutant pair YH, without a change in the rotamer states. It is also possible that AF2 predicted rotamer states are incorrect in some cases, falsely predicting no rotamer state changes pairs that are cooperative. Experimental structures would be needed to test this. For the pair V422Q/P484F (ABL1), although no experimental structure of the double mutant is available, an experimental structure is available for a different kinase protein with the QF residue pair at position corresponding to 422/484 in the MSA, tuberculosis PknI kinase, and it shows a different rotamer state than the AF2 prediction of the QF double mutant, suggesting a possible AF2 rotamer misprediction. The remaining 5 cooperative double mutant pairs for PIM1 exhibit similar behavior to those of ABL1.

**Table 1.**
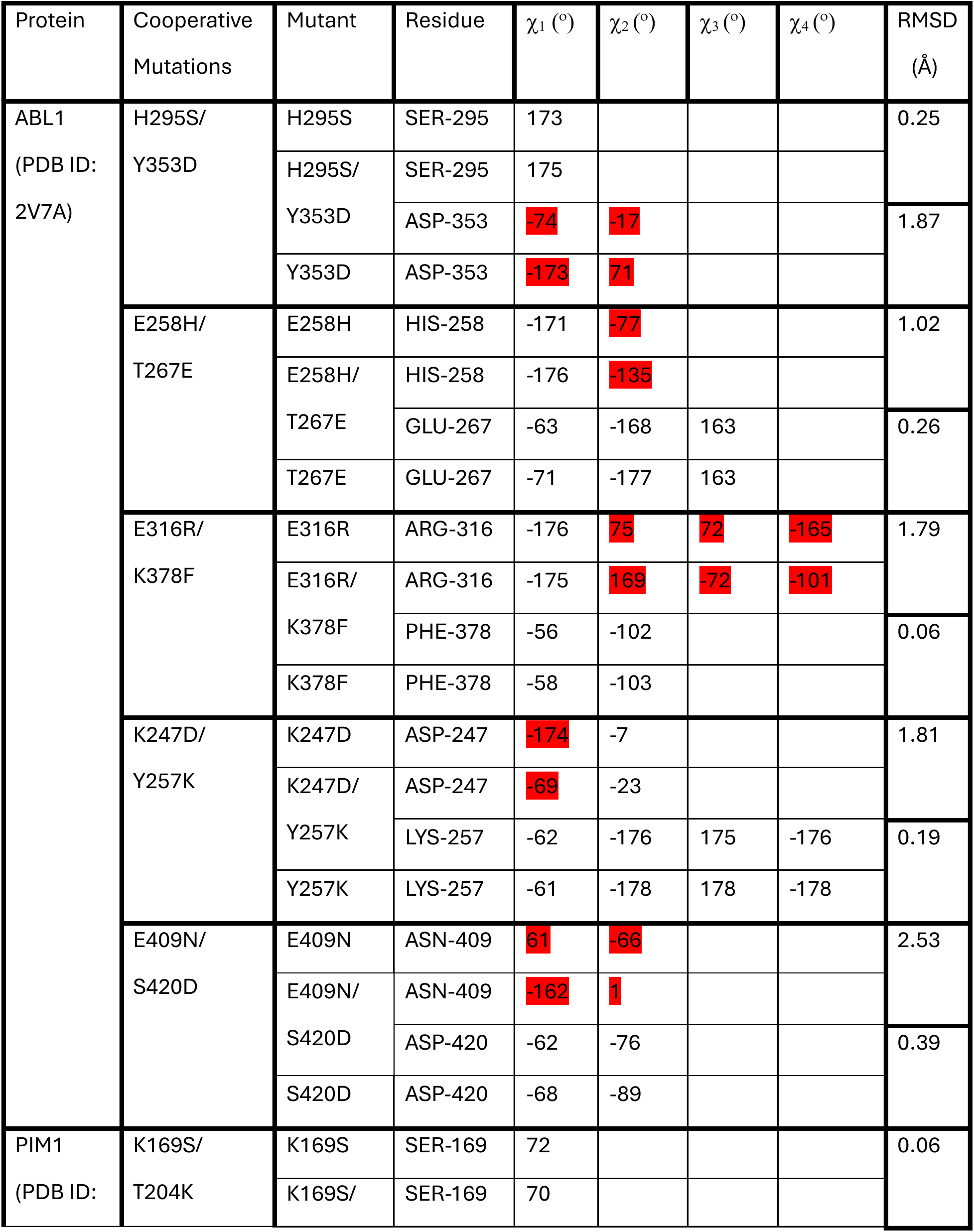
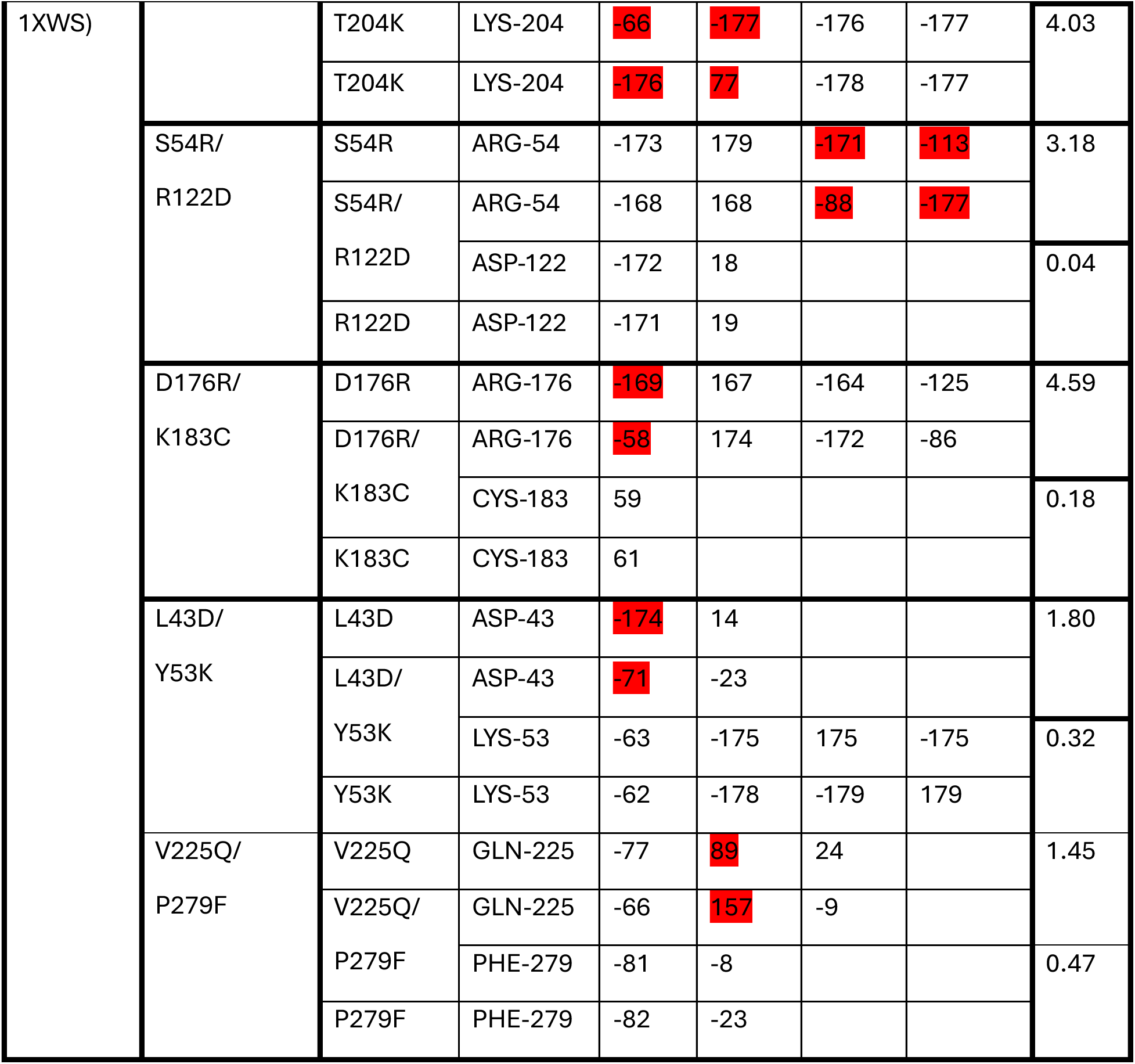
Mutant pairs with the double mutant rotamer states differs from the single mutant rotamer states with differences highlighted in red.

In this section we have illustrated how the Potts sequence-based energy model can be used to perform mutational scans of a protein to search for the most strongly cooperative mutational pairs, and AlphaFold can then be used to detect the structural changes associated with the very strong cooperativity. Further developing these ideas, it should be possible to integrate the Potts model and AlphaFold into a single pipeline which can search for mutational pairs that have the strongest effects on protein fitness, build structural models of these mutant proteins and their assemblies, and use these models to guide experiments designed to probe the role this cooperativity plays in the protein’s function and dysfunction.

## Conclusions

We have analyzed the side chain rotamer states predicted by ColabFold for every amino acid residue in ten different proteins; a total of 1453 side chain rotamer state predictions. ColabFold is an online implementation of AlphaFold2. Unlike the striking accuracy (the root mean square deviation of C^a^ atoms averaged over the ten studied proteins is less than 1Å) observed between backbones of experimental PDB and ColabFold predicted structures, discrepancies between side-chain angles of experimental and predicted structures are significantly larger. Using a deviation of more than +/− 40 degrees from the experimental value as a definition of side chain error, we find that over a set of 10 benchmark proteins, the rotamer state prediction error of ColabFold is on average ∼14% for χ_1_ dihedral angles, and increases to ∼48% for χ_3_ dihedral angles. The prediction error is smaller for non-polar side chains and is improved using structural templates. When structural templates are employed, the largest improvement is for χ_1_ dihedrals (∼31% improvement compared to without templates).

ColabFold demonstrates a bias towards the most prevalent rotamer states in the protein data bank, potentially limiting its ability to capture rare side chain conformations effectively. The rotamer state prediction error rate increases near Alanine and Glycine residues, which surprisingly appears to be caused by a bias for ColabFold to predict the most populated rotamer states in the protein data bank, while the lack of packing constraints adjacent to Alanine and Glycine residues or the increased backbone flexibility near these residues may play a less significant role. We hope these observations can lead to approaches to improve the training of AlphaFold to better account for rotamer states of residues that are only rarely observed in the PDB.

As a first application of AlphaFold to explore the structural consequences of strongly cooperative mutations on side chain rearrangements, we employ a Potts sequence-based energy model to perform large scale mutational scans of two proteins ABL1 and PIM1 kinase, searching for the most strongly cooperative mutational pairs, and then use ColabFold to predict the structural signatures of this cooperativity on the interacting side-chains. The predictions of the sequence-based Potts statistical energy model, and structure based AlphaFold are largely consistent, in that for more than 50% of the examples, the structure of one side chain mutant significantly perturbs the structure of the second side chain (either wild type or mutant), leading to increased stability of the folded protein according to the Potts sequence-based statistical energy model; in the remaining cases the increased stability can be rationalized without requiring a rotamer state change. Our results demonstrate that the integration of the sequence-based Potts model with AlphaFold into a single pipeline provides a new tool that can be used to explore the fundamental relationship between protein mutations, cooperative changes in structure, and fitness.

## Appendix: Introduction to the Potts model Hamiltonian and derivation of equation 1 in the main text

The cooperative mutation effects can be modeled using statistical mechanics and Boltzmann networks by learning the mutational patterns in the protein families using Multiple Sequence Alignments (*MSA*) (57). Our sequence-based double mutant cooperativity (ΔΔ*E*) calculation is based on the *MSA* used to derive the protein Kinase Potts Hamiltonian model which describes the coevolutionary interactions between all possible residue pairs in the sequence with all possible amino acid combinations at each pair of positions. Analogous to the Hamiltonian of the magnetic spin system in the Condensed Matter Physics, it is defined by,

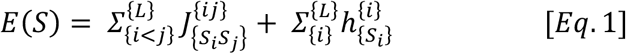

where 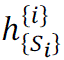 and 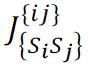 are the fields and couplings, also called single-site and pairwise statistical energy terms, respectively (58). The coupling term gives the strength of the interaction between the residues *S_i_* and *S_j_* at positions i and j, respectively whereas 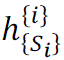 represents the field for residue Si at position i in the sequence *S*. The indices i and j represent the amino acid positions in the aligned sequence *S* of length *L*. Each position can have *20* possible states (*q=20*) with one gap character. The statistical energy E of a sequence of length *L* gives the probability of that sequence to be observed in the MSA relative to all other sequences following the Boltzmann distribution.

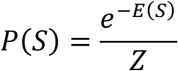

where 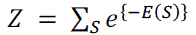 is the partition function that takes sum over all possible sequences and ensures the normalization.

The model is parametrized such that it exactly reproduces the following *MSA* marginals, the single-site residue frequencies and joint frequencies of every pair of residues in the *MSA*, also referred to as univariate and bivariate residue frequencies, respectively, and are defined by:

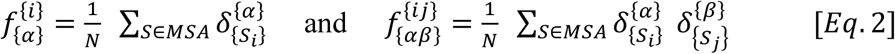

Here, *N* is the *MSA* depth (number of sequences in the *MSA*), and δ is Kronecker delta function where 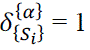 if the residue α is observed at position i in the sequence *S*, and 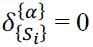 otherwise.

The univariate and bivariate marginals from the model is difficult to estimate directly as it requires computing partition function *Z* which takes sums over *q*^*l*^ sequences which is complex. There are several methods for inferring the model parameters such that the model distribution *P(S)* reproduces the empirical univariate and bivariate marginals of the *MSA*. The protein Kinase Potts Hamiltonian Model implements Markov Chain Monte Carlo (MCMC) based Mi3-GPU method for such purpose. Through the MCMC sampling, it generates a synthetic *MSA* from which marginals can be calculated. With an initial guess for the couplings, the model generates synthetic *MSA* from which marginals are calculated and used those values to iteratively update the coupling parameter until the empirical marginals of the *MSA* dataset are recovered. The resulting Hamiltonian can generate the sequences whose univariate and bivariate marginals match with empirical *MSA* marginals. The detailed explanation can be studied in (59).

The Potts model can be used to estimate the cooperative effect of a pair of mutations on protein fitness. This double mutant cooperativity is defined by subtracting the additive effects of two single mutations at position i and j from the double mutation effect. Mathematically, it is defined by:

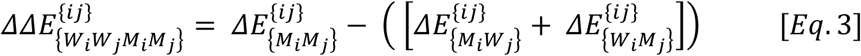

The first term in right hand side gives the change in energy when both residues i and j are mutated while the two terms inside parenthesis contribute energy change due to single mutation at position *i* and *j*, respectively. These terms can be further defined as:

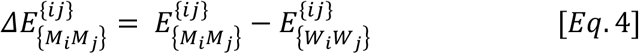

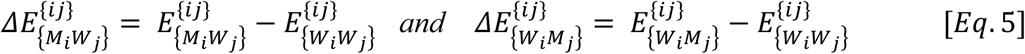

Substituting [*Eq*. 4] and [*Eq*. 5] in [*Eq*. 3], we get:

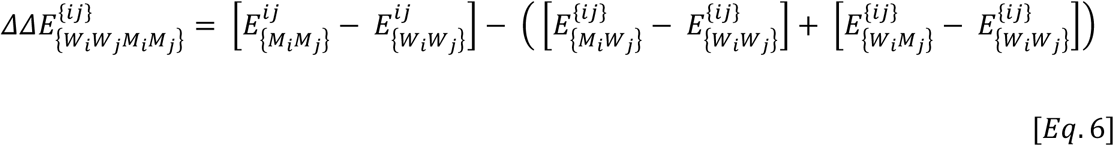

where the first term in right hand side represents the relative log probability of the double mutation *(M_i_, M_j_)* at positions i and j relative to the reference residues *(W_i_,W_j_)*, while the second and third terms are the corresponding log relative probabilities of the two single mutations. After using *[Eq.1]* in [*Eq.6*], the field contributions to each of the terms cancel when combined in the ΔΔ*E* formula, and *[Eq.6]* can be written:

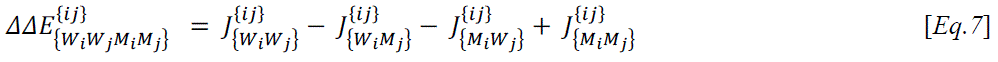

ΔΔ*E* then depends on the four couplings between the residues pairs *(W_i_,W_j_), (M_i_, M_j_), (W_i_, M_j_)* and (*M_i_, W_j_*) at positions *i* and *j*. Importantly, ΔΔ*E* is independent of the gauge of the Potts statistical energy function. The non-linear cooperativity is a measure of how much more favorable (or unfavorable) the fitness of the double mutation is than the sum of the effects of the individual mutations on the fitness.

### Author Contributions

G.G.M. and R.M.L. designed research, G.G.M., A.T., K.K., and A.H. performed research, G.G.M., A.T., and R.M.L. analyzed data, G.G.M., A.T., K.K., A.H., and R.M.L. wrote the manuscript.

### Declaration of Interests

The authors declare no competing interests.

## Acknowledgement

We thank Dr. Sompriya Chatterjee for performing the RMSD calculations and providing the scripts used to perform these calculations. This work was supported in part by the National Institutes of Health grant NIH R35 GM132090.

