## Supplemental Figures (S1-S6) and Tables (S1, S2) for "Using AlphaFold2 to Predict the Conformations of Side Chains in Folded Proteins"

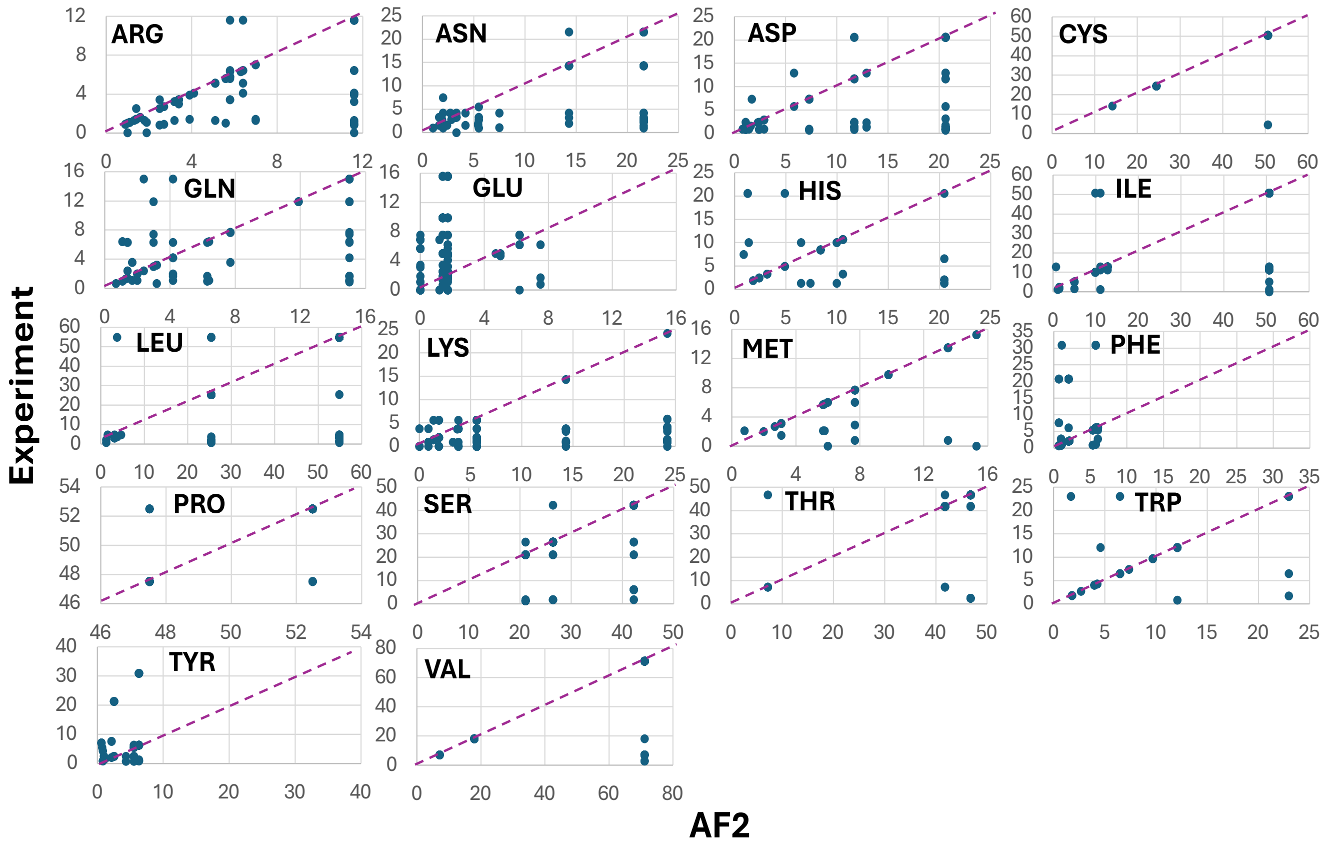


Figure S1. Comparison of rotamer states of each amino acid of experimental and AF2 predicted (with template) structures in terms of the probability (in %) of the closest rotamer state in the protein database.

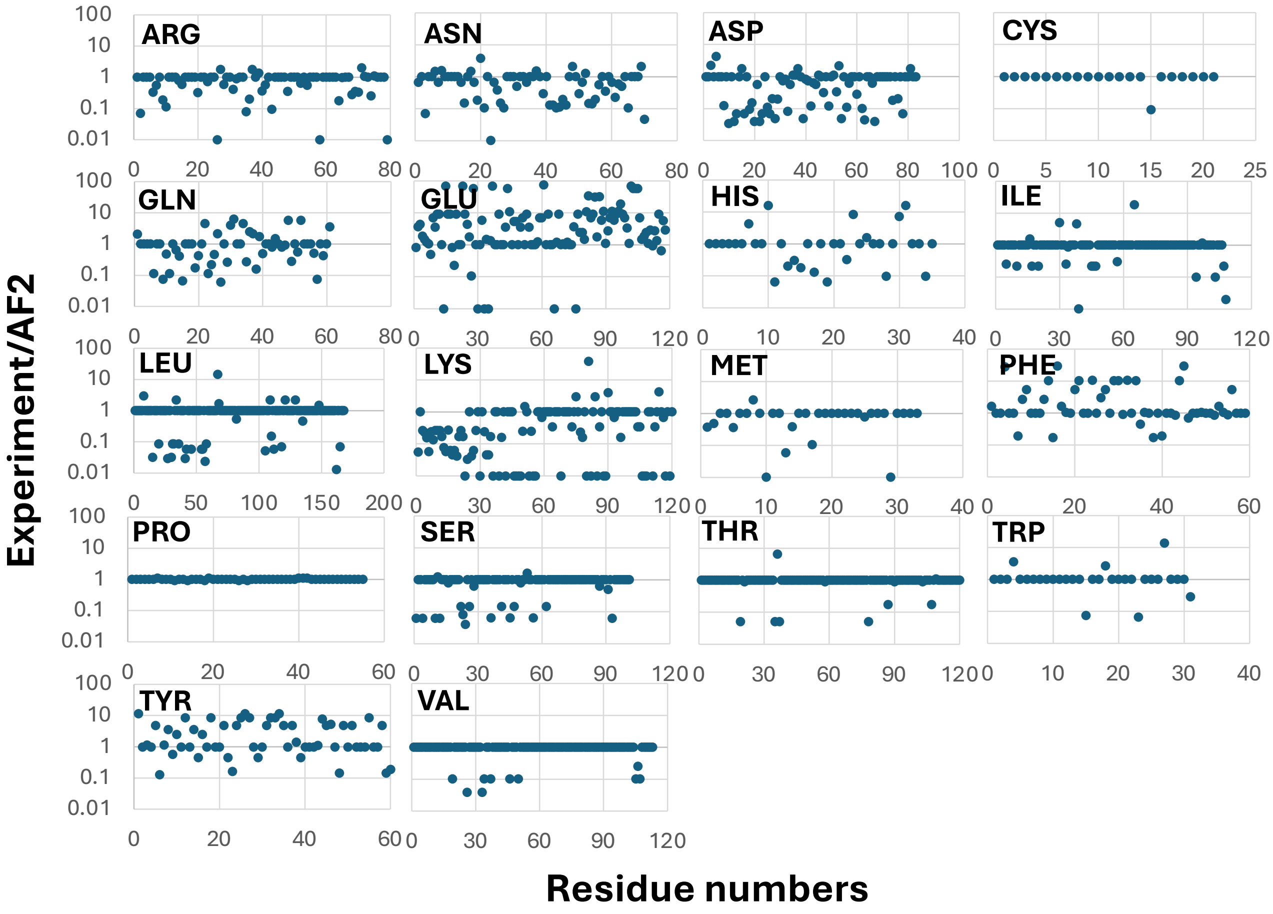


Figure S2. The ratios between the PDB derived database percentages for each residue of ten experimental and AF2 predicted structures. If a ratio is less than 1, it indicates that a rotamer state of that particular amino acid in AF2 predicted structure is inclined towards dominant rotamer states in protein data base compared to its counterpart in experimental structure. If a ratio is more than 1, it indicates that a rotamer state of that particular amino acid in experimental structure tends to adopt the dominant conformation from protein data base compare to its counterpart in AF2 predicted structure. If a ratio is equal 1, it indicates that a rotamer states of that particular amino acid in AF2 predicted and experimental structures do not differ from each other and fall into the same category defined by protein data base.

### Non-polar Amino Acids

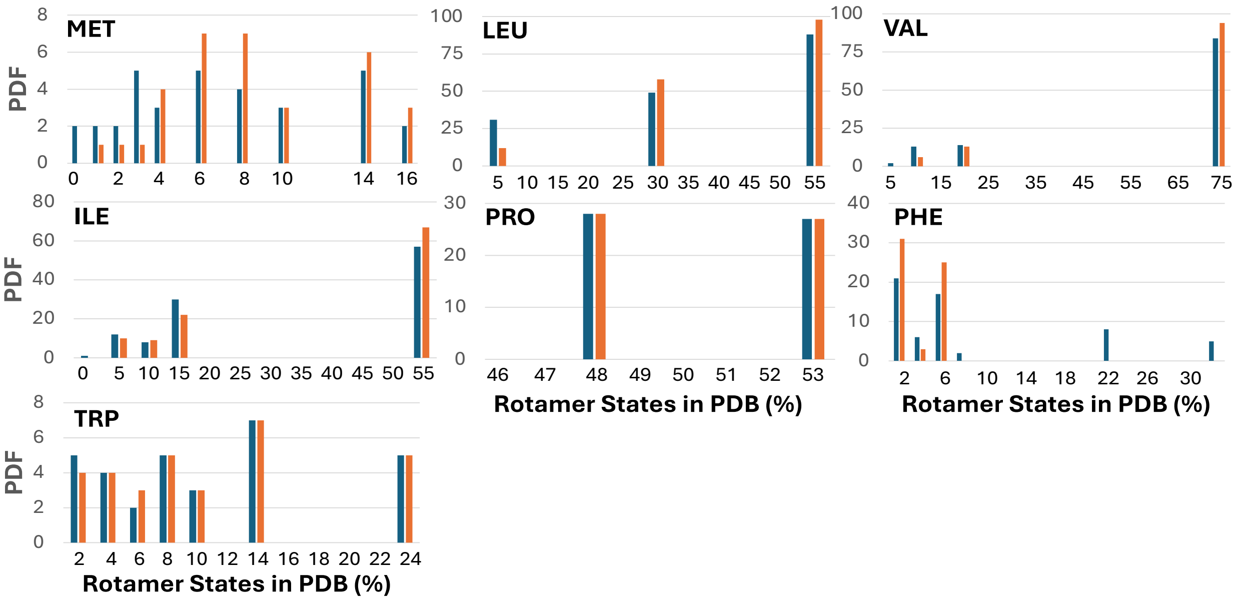


### Polar Uncharged Amino Acids

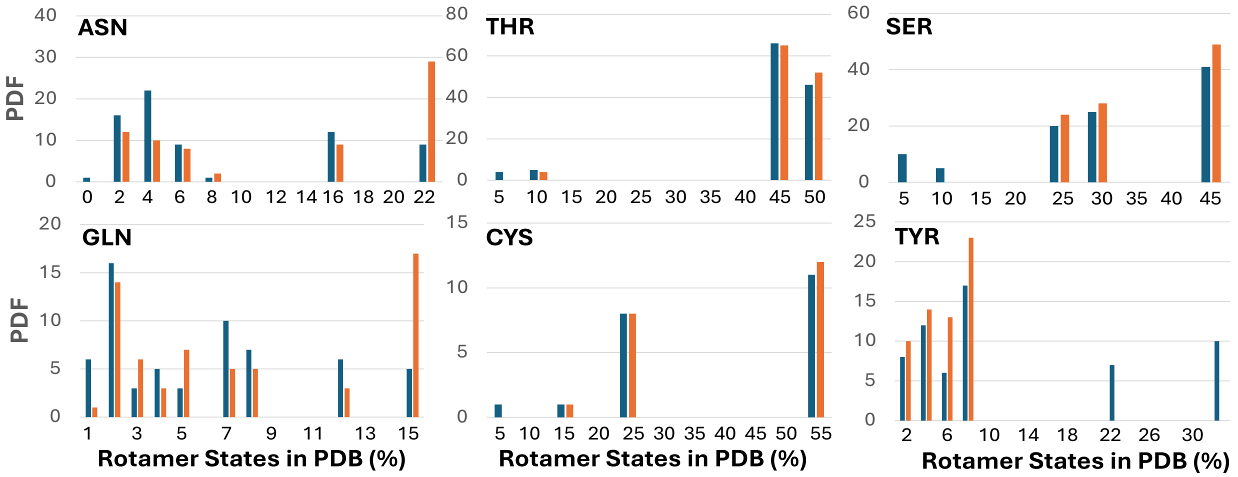


### Polar Charged Amino Acids

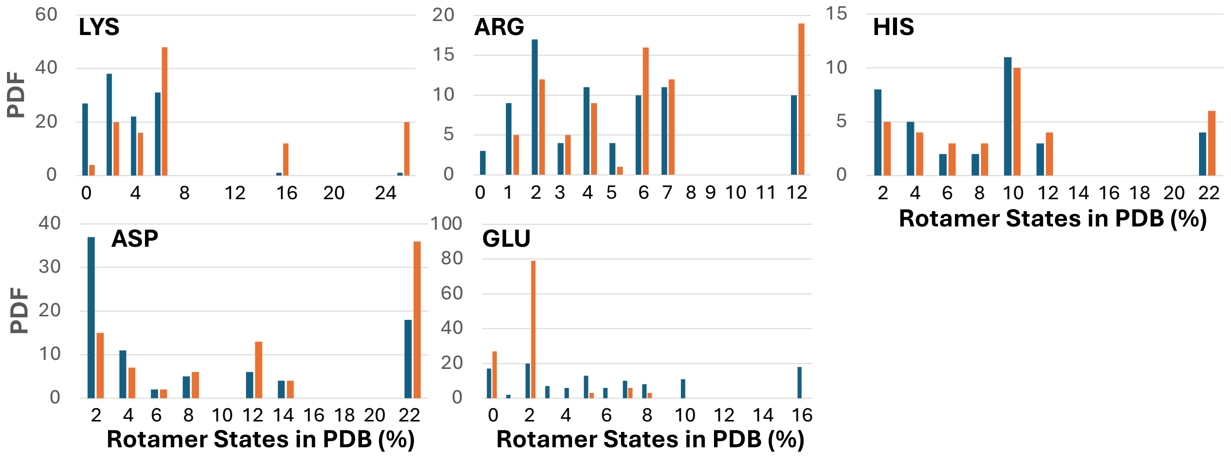

Figure S3. Probability distribution functions of rotamer states of amino acids obtained from ten experimental (blue bars) and AF2 predicted (orange bars) structures along the populations of rotamer states in protein data base.

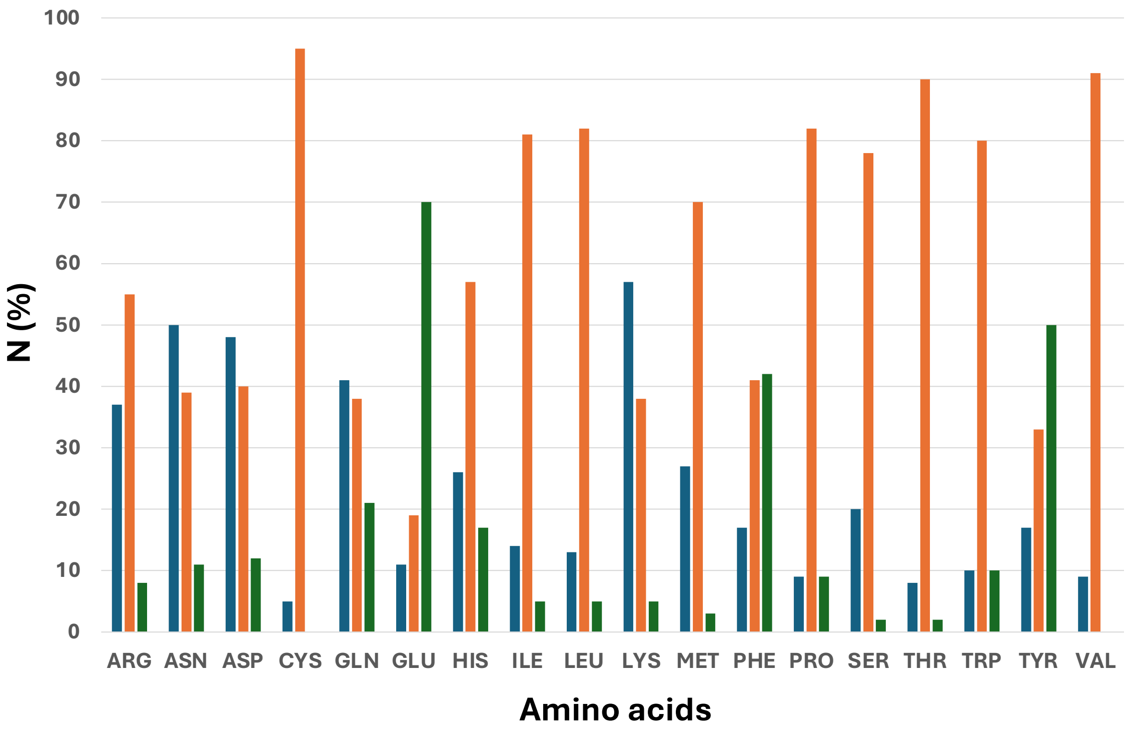


Figure S4. Distribution of percentages of how often the rotamer states of each amino acid in AF2 predicted (blue bars) and experimental (green bars) structures adopt more dominant rotamer state from protein data base, and how often they do not differ from each other (orange bars).

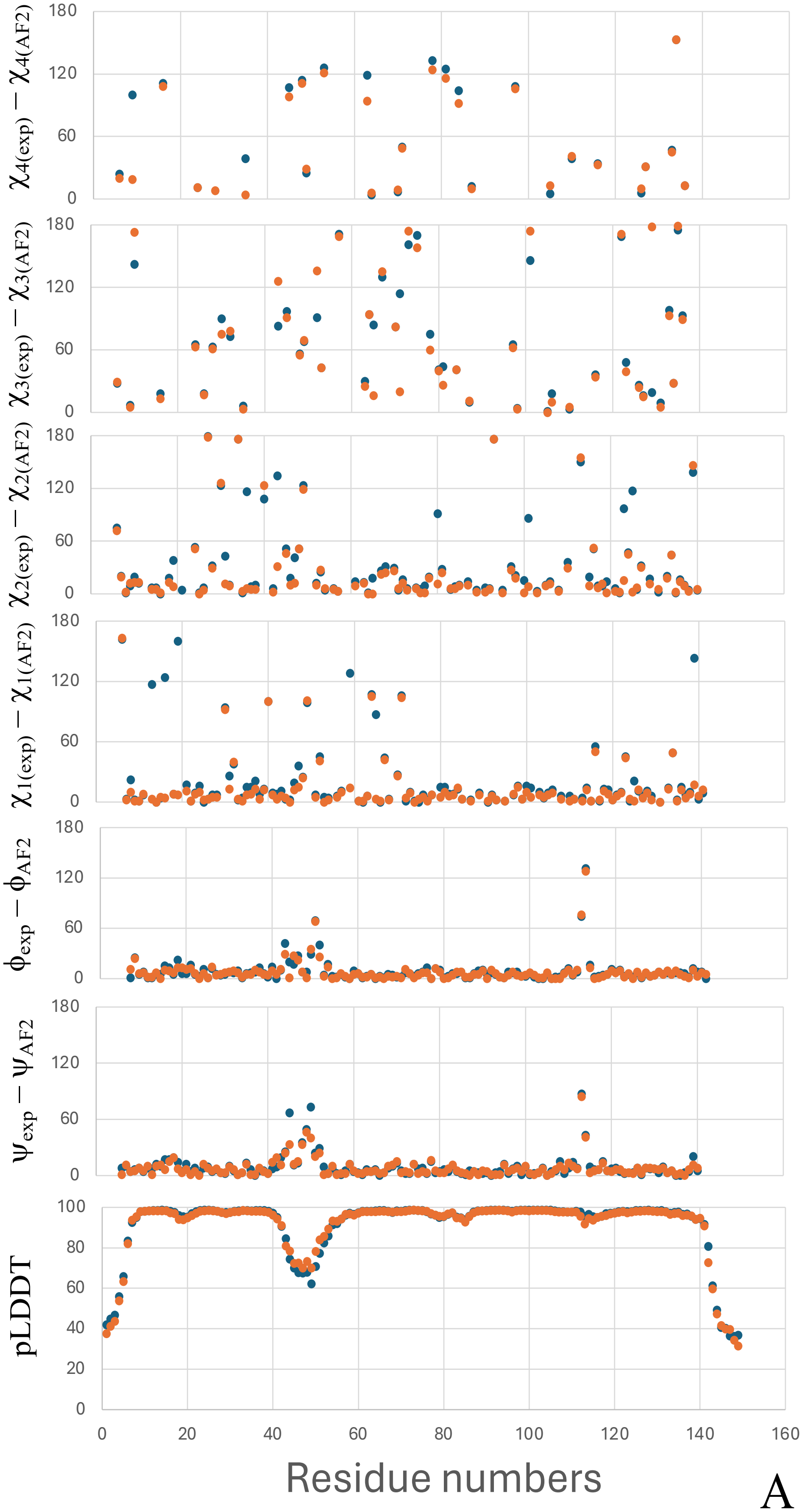

180
120
60
0
$\chi_{4(exp)} - \chi_{4(AF2)}$
$\chi_{3(exp)} - \chi_{3(AF2)}$
$\chi_{2(exp)} - \chi_{2(AF2)}$
$\chi_{1(exp)} - \chi_{1(AF2)}$
$\phi_{exp} - \phi_{AF2}$
$\psi_{exp} - \psi_{AF2}$
pLDDT
100
80
40
20
0 20 40 60 80 100 120 140 160
Residue numbers


A

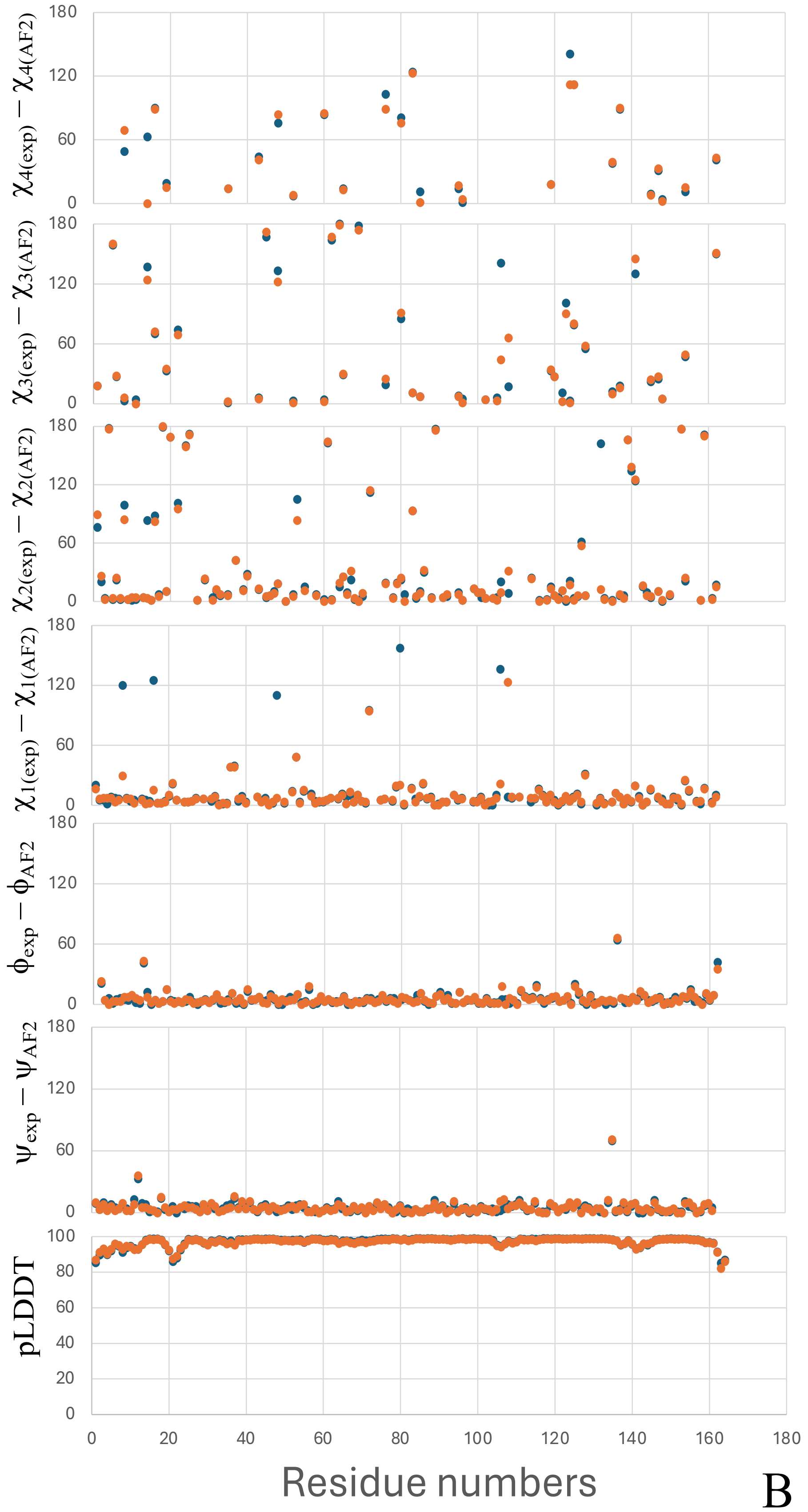
$\chi_{4(exp)} - \chi_{4(AF2)}$
$\chi_{3(exp)} - \chi_{3(AF2)}$
$\chi_{2(exp)} - \chi_{2(AF2)}$
$\chi_{1(exp)} - \chi_{1(AF2)}$
$\phi_{exp} - \phi_{AF2}$
$\psi_{exp} - \psi_{AF2}$
pLDDT
Residue numbers

B

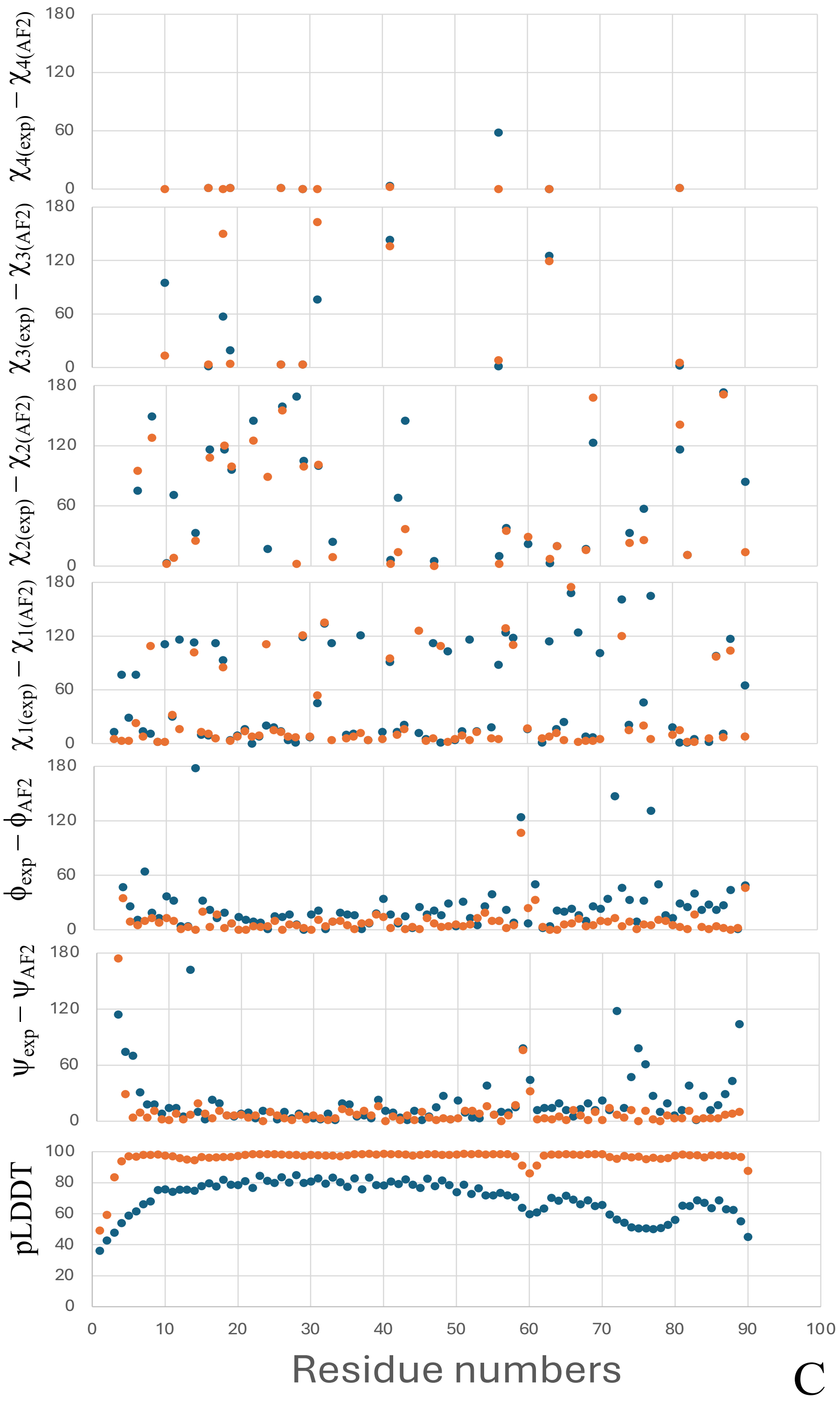


Figure S5. The pLDDT scores, the errors between the values of backbone (ψ,ϕ) and side-chain ($\chi_1$, $\chi_2$, $\chi_3$, $\chi_4$) angles for experimental and AF2 predicted structures with (orange dots) and without (blue dots) template are illustrated for 1EY0 (panel A), 1L63 (panel B) and 1L0S (panel C) proteins.

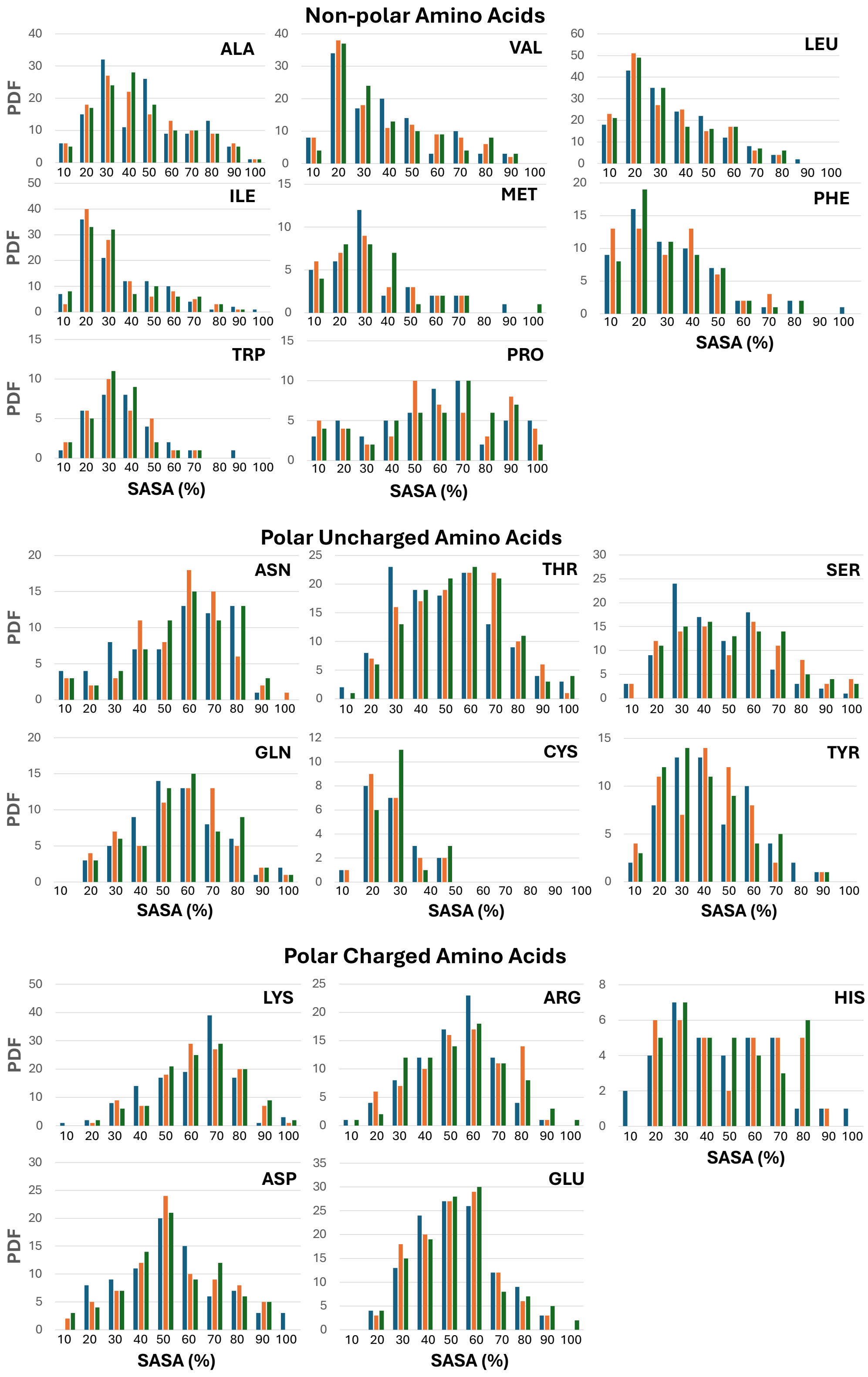
Non-polar Amino Acids
ALA
VAL
LEU
ILE
MET
PHE
TRP
PRO
Polar Uncharged Amino Acids
ASN
THR
SER
GLN
CYS
TYR
Polar Charged Amino Acids
LYS
ARG
HIS
ASP
GLU
PDF
SASA (%)

Figure S6. Probability distribution functions (PDFs) of percentages of SASA for non-polar, polar uncharged and polar charged amino acids computed from the experimental (blue bars) and predicted structures with- (orange bars) and without (green bars) templates for 1EY0, 1L0S, 1L63, 1XWS, 2LZM, 2V7A, 3MSW, 3V2V, 3VN3, and 6B87 proteins.

Table S1. Rotamer state populations (five the most dominant rotamer states) for amino acids in the Schrödinger knowledge base

| Amino Acid | 1st (%) | 2nd (%) | 3rd (%) | 4th (%) | 5th (%) |
|---|---|---|---|---|---|
| Arg | 11.6 | 7.0 | 6.4 | 6.3 | 5.8 |
| Asn | 21.6 | 14.3 | 7.5 | 5.5 | 5.4 |
| Asp | 20.6 | 12.9 | 11.7 | 7.3 | 5.8 |
| Cys | 50.6 | 24.5 | 14.2 | 4.6 | 2.4 |
| Gln | 15.0 | 11.9 | 7.7 | 7.4 | 6.4 |
| Glu | 15.6 | 9.9 | 7.5 | 6.9 | 6.2 |
| His | 20.5 | 10.6 | 10.0 | 8.4 | 7.4 |
| Ile | 50.6 | 12.8 | 11.1 | 9.9 | 5.0 |
| Leu | 54.9 | 25.4 | 4.7 | 3.8 | 3.2 |
| Lys | 24.2 | 14.3 | 7.4 | 5.8 | 5.6 |
| Met | 15.3 | 13.5 | 9.8 | 7.7 | 6.0 |
| Phe | 30.8 | 20.7 | 7.5 | 7.2 | 6 |
| Ser | 42.2 | 26.4 | 21.1 | 6.1 | 1.7 |
| Thr | 46.8 | 41.8 | 7.2 | 2.4 | 1.0 |
| Trp | 23.0 | 12.1 | 9.7 | 7.4 | 6.5 |
| Tyr | 30.9 | 21.2 | 7.6 | 7.0 | 6.3 |
| Val | 71.2 | 18.0 | 7.1 | 2.7 | 0.9 |
| Pro* | 52.5 | 47.5 | | | |

*Data are obtained from http://dunbrack.fccc.edu/lab/conformational_analysis#Intro

Table S2. Correlations between errors in backbone RMSD values and side chain rotamer states.

| Proteins | Pearson's correlation coefficient between RMSD and $\chi_1$ | Average RMSD (in Å) for ALA residues | Average RMSD (in Å) for GLY residues | Average RMSD (in Å) for entire sequence without ALA and GLY | Average RMSD (in Å) for residues with large $\Delta\chi_1$ ($\chi_{1exp} - \chi_{1AF2} > 40°$) | Large backbone RMSD errors (>0.5Å) (in %) at loops and ends | Large backbone RMSD errors (>0.5Å) (in %) at α-helices and β-sheets |
|---|---|---|---|---|---|---|---|
| 1EY0 | 0.21 | 0.32 | 0.36 | 0.35 | 0.48 | 100 | 0 |
| 1L63 | -0.04 | 0.36 | 0.31 | 0.38 | 0.31 | 72 | 28 |
| 1L0S | 0.01 | 0.29 | 0.51 | 0.25 | 0.24 | 57 | 43 |
| 2LZM | 0.06 | 0.44 | 0.45 | 0.50 | 0.62 | 53 | 47 |
| 2V7A | 0.21 | 0.28 | 0.70 | 0.32 | 0.44 | 73 | 27 |
| 1XWS | 0.26 | 0.13 | 0.35 | 0.21 | 0.41 | 100 | 0 |
| 3V2V | 0.19 | 0.64 | 0.55 | 0.55 | 0.71 | 33 | 67 |
| 6B87 | 0.22 | 0.29 | 0.30 | 0.34 | 0.48 | 36 | 64 |
| 3VN3 | 0.28 | 0.18 | 0.21 | 0.19 | 0.34 | 100 | 0 |
| 3MSW | 0.10 | 1.33 | 0.60 | 0.35 | 0.56 | 100 | 0 |
